# Learning context-aware, distributed gene representations in spatial transcriptomics with SpaCEX

**DOI:** 10.1101/2024.06.07.598026

**Authors:** Xiaobo Sun, Yucheng Xu, Wenlin Li, Mengqian Huang, Ziyi Wang, Jing Chen, Hao Wu

**Author notes:** X.S. and H.W. conceived and supervised the study. X.S. derived the model and developed the framework. X.S., H.W., W.L. wrote and revised the manuscript. X.S., Y.X., W.L., and Z.W. implemented the framework. Y.X., W.L., M.H., and J.C. conducted the experiments. X.S., Y.X., M.H., Z.W. conducted the analyses. Y.X., W.L., X.S., M.H., and J.C. collected the results and plot the figures. All authors approved the manuscript. The authors declare no competing interests.

## Abstract

Distributed gene representations are pivotal in data-driven genomic research, offering a structured way to understand the complexities of genomic data and providing foundation for various data analysis tasks. Current gene representation learning methods demand costly pretraining on heterogeneous transcriptomic corpora, making them less approachable and prone to over-generalization. For spatial transcriptomics (ST), there is a plethora of methods for learning spot embeddings but serious lacking method for generating gene embeddings from spatial gene profiles. In response, we present SpaCEX, a pioneer cost-effective self-supervised learning model that generates gene embeddings from ST data through exploiting spatial genomic “context” identified as spatially co-expressed gene groups. SpaCEX-generated gene embeddings (SGE) feature in context-awareness, rich semantics, and robustness to cross-sample technical artifacts. Extensive real data analyses reveal biological relevance of SpaCEX-identified genomic contexts and validate functional and relational semantics of SGEs. We further develop a suite of SGE-based computational methods for a range of key downstream objectives: identifying disease-associated genes and gene-gene interactions, pinpointing genes with designated spatial expression patterns, enhancing transcriptomic coverage of FISH-based ST, detecting spatially variable genes, and improving spatial clustering. Extensive real data results demonstrate these methods’ superior performance, thereby affirming the potential of SGEs in facilitating various analytical task.

**Significance Statement:** Spatial transcriptomics enables the identification of spatial gene relationships within tissues, providing semantically rich genomic “contexts” for understanding functional interconnections among genes. SpaCEX marks the first endeavor to effectively harnesses these contexts to yield biologically relevant distributed gene representations. These representations serve as a powerful tool to greatly facilitate the exploration of the genetic mechanisms behind phenotypes and diseases, as exemplified by their utility in key downstream analytical tasks in biomedical research, including identifying disease-associated genes and gene interactions, *in silico* expanding the transcriptomic coverage of low-throughput, high-resolution ST technologies, pinpointing diverse spatial gene expression patterns (co-expression, spatially variable pattern, and patterns with specific expression levels across tissue domains), and enhancing tissue domain discovery.

---

**D**istributed gene representations embed multifaceted nature of genes within a high-dimensional space, offering profound insights into the complex mechanisms of gene expression, regulation, and interaction, and paving ways for leveraging machine learning techniques in advancing biomedical research (1), disease diagnosis (2), and the discovery of therapeutic targets (3) with unprecedented precision and efficiency.

Spatial transcriptomics (ST), including high-resolution in situ hybridization-based (e.g., SeqFISH (4)) and high-throughput in situ capturing-based (e.g., 10x Visium (5)) technologies, enables the profiling of spatial gene expression in heterogeneous tissues, providing unprecedent opportunities to characterize spatial distribution of cell types (6), delineate spatial tissue organization (5), gene-gene interactions (7), etc. ST also can reveal spatial genomic “contexts” formed by genes cofunctional in the same biological processes and pathways, given the similarity of these genes in spatial expression patterns within tissues. Such genomic contexts not only provide insights into the molecular mechanisms underlying biological processes and disease development (8), but also allow the learning of distributed gene representations, resembling the learning of word representations from word contexts in linguistic models (1, 9, 10). These gene embeddings provide a foundation for quantitatively characterizing context-specific gene functions and interactions from a spatial perspective, facilitating various analytical endeavors where insights into spatial genetic mechanisms are critical.

However, to the best of our knowledge, there is currently no method for learning gene embeddings from ST data due to challenges in effectively identifying spatial genomic contexts and encoding spatial gene expression patterns. While recent works, including Gene2vec (1), scGPT (9), scFoundation (11), scBERT (12), and geneFormer (13), have been developed to learn gene embeddings from atlas-scale microarray or scRNA-seq data, they do not extend to ST, overlooking crucial spatial gene expression information. Moreover, it has been observed that the extensive pretraining of these models on massive data corpora offers marginal benefits for finetuning downstream tasks (14). This is probably due to the irreconcilable heterogeneities in pretraining data collected across a variety of technical and biological conditions, which hinder the model’s ability to grasp context-specific gene embedding nuances. In addition, these methods, characterized by their tremendous amount of parameters, are prone to sensitivity regarding hyper-parameters settings and parameter initializations (14). This sensitivity necessitates repeated pretraining, particularly with the introduction of new data, which, coupled with the prohibitive costs of data collection and model pretraining, renders the updating of pretrained models impractical for the broader research community.

To bridge this gap, we develop a “context-aware, self-supervised learning on **Spa**tially **C**o-**EX**pressed genes (SpaCEX)” model. SpaCEX features in utilizing spatial genomic context inherent in ST data to generate gene embeddings that accurately represent the condition-specific spatial functional and relational semantics of genes. Technically, SpaCEX treats gene spatial expressions as images and leverages a masked-image model (MIM), which excels in extracting local-context perceptible and holistic visual features (15), to yield initial gene embeddings. These embeddings are iteratively refined through a self-paced pretext task aimed at discerning genomic contexts by contrasting spatial expression patterns among genes, drawing genes with similar spatial expressions closer in the latent embedding space, while distancing those with divergent patterns. This step enhances the discriminability of gene embeddings, facilitating their incorporation of intricate relational semantic structures among genes. The innovative integration of MIM with contrastive learning equips SpaCEX with a superior capability to comprehend and translate spatial gene information, including both expression patterns and inter-gene relationships, into **S**paCEX-generated **G**ene **E**mbeddings (SGE).

Our study is organized as follows: Firstly, to establish the validity of our method, we demonstrate SpaCEX’s effectiveness in identifying spatially co-expressed gene groups, justifying their role as genomic contexts by showing their biological relevance. In addition, the functional and relational semantics of SGEs undergo rigorous validations. Particularly, we demonstrate SGEs’ robustness to technical variations and their utility in gene alignment and comparison across samples. Secondly, we demonstrate the applicability of SpaCEX and SGEs in key downstream tasks by developing a suite of SGE-based computational methods for: i) the identification of disease-associated genes and gene crosstalk; ii) the de novo expansion of transcriptomic coverage of FISH-based ST data, addressing a longstanding challenge that has restricted the broader application of FISH-based ST data; iii) pinpointing genes with designated spatial expression patterns in tissues; iv) detecting spatially variable genes (SVGs); v) improving spatial clustering. Extensive real data analyses demonstrate that our methods either provide optimal solutions to challenges that have been not inadequately addressed, e.g., the first three tasks, or outperform established benchmarks as seen in task iv and v. These tasks exemplify how SGEs can be effectively employed to address various downstream task-specific objectives, promising SpaCEX’s potential in developing a genomic “language”-based methodological ecosystem.

## Result

### Overview of SpaCEX

The fundamental idea (Fig. 1*A*) of SpaCEX is that genes co-functional in gene networks and pathways typically exhibit similar spatial expression patterns in tissues, forming a “genomic context” resembling the word context in natural languages. Through a self-supervised learning of the “proximity” of genes in spatial transcriptional activity, as implied by the ST data, we can concurrently identify spatial genomic contexts and acquire semantically meaningful gene embeddings in a data-driven manner. These embeddings, representing spatial gene functions and relationships, can facilitate downstream task-specific objectives. As illustrated in Methods and Fig. 1*B*, SpaCEX mainly consists of three modules. In *Module I*, erage an adapted masked autoencoder (MAE) (15) to tranansform gene images into gene embeddings that follow a mixture of multivariate Student’s t distributions (see *“Representation learning of spatial gene expression maps”* in Methods). With this MIM, SpaCEX gains the local-context perceptibility by learning to regenerate masked image patches from the surrounding contexts. In *Module II*, the gene embeddings are modeled as a Students’ t mixture model (SMM) in the latent feature space, with each mixture component serving as a genomic context comprising spatially co-expressed genes. After the estimation of SMM parameters via a Maximum a *posterior* (MAP)-EM algorithm, soft assignments of genes to the mixture components are computed (see *“SMM-based modeling”* in Methods). *Module III* implements a semi-contrastive learning process, through which SpaCEX gains the ability to discriminate between spatial gene expressions and grasps the intricate relational semantic structures among genes. This process involves a self-paced, iterative joint optimization of MAE weights and SMM parameters using two loss functions *ℒ* _1_ and *ℒ*_2_ (see *“Self-paced semi-contrastive optimization of gene embeddings”* in Methods). Each training epoch begins with *ℒ*_1_, updating the MAE weights to maximize the log likelihood of the entire dataset while controlling for macro-factors (e.g., cluster size imbalance) via regularization terms. Following this, *ℒ*_2_, designed for discriminatively boosted clustering, further refine gene embeddings and SMM parameters, drawing closer similar genes and distancing dissimilar ones over successive batches. Overall, the training process alternates between the *Module II* and *III* until either a predetermined number of training epochs is reached, or the change in gene assignments between successive epochs falls beneath a prespecified threshold.

**Fig. 1.**
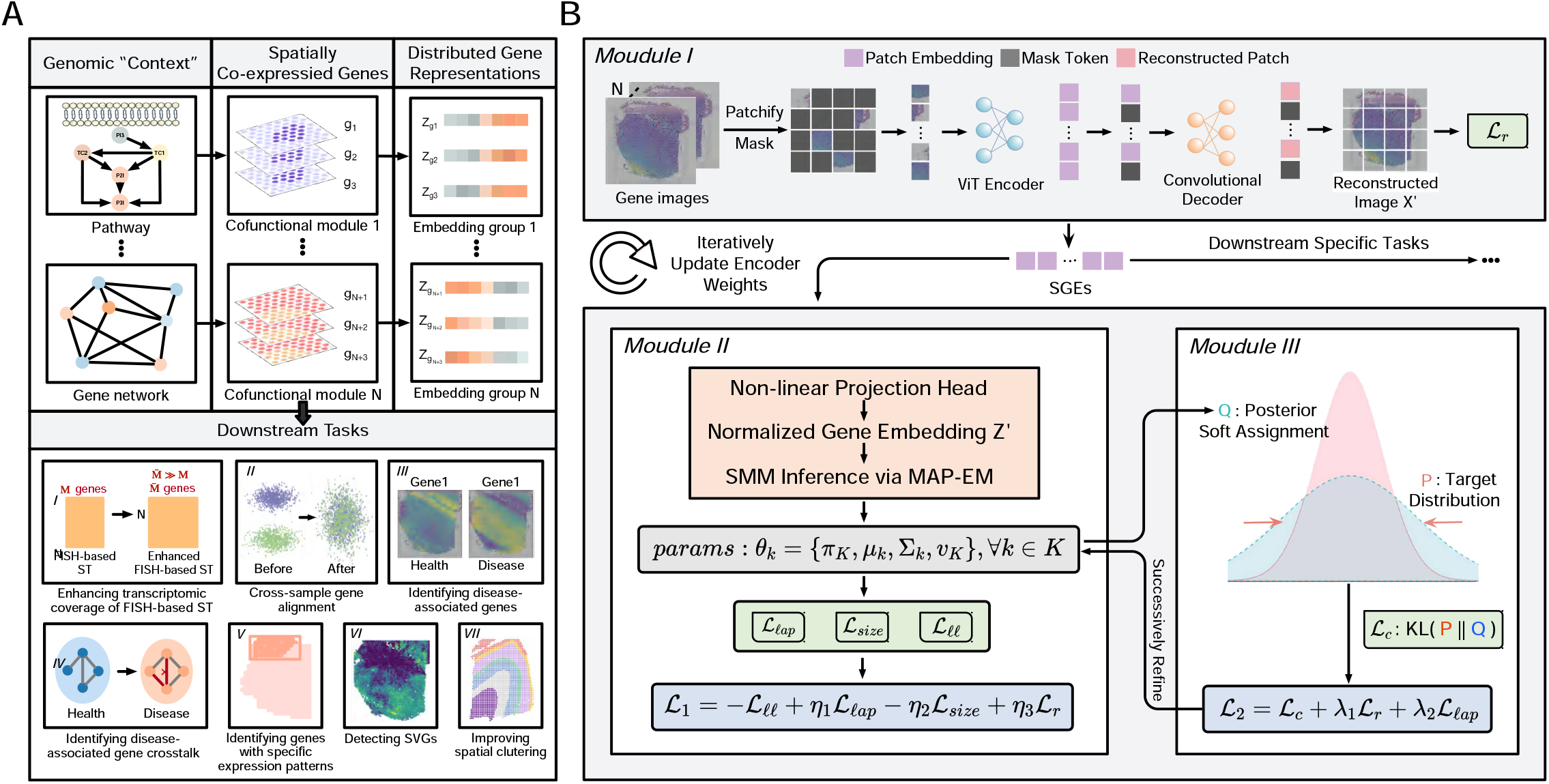
Overview of the distributed gene representation learning with SpaCEX. (*A*) Outline of learning distributed gene representations from spatial genomic contexts. ST data reveal spatial genomic contexts comprising genes cofunctional in gene pathways and networks since they tend to exhibit similar expression patterns across tissue space. By leveraging these genomic contexts, distributed gene representations can be learned. These representations not only encapsulate gene spatial functional and relational semantics but also instrumental in facilitating various downstream task-specific objectives. (*B*) Workflow of SpaCEX. It comprises three modules: In Module I, SpaCEX employs an adapted masked autoencoder (MAE) to learn the representations of gene images, which enhances gene embeddings’ local-context perceptibility. Module II involves modeling gene embeddings using a Student’s t mixture model, with parameters estimated via a MAP-EM algorithm. This module aims to maximize the likelihood of the entire dataset. In *Module III*, gene embeddings are refined through a self-paced pretext task that aims to identify genomic contexts via iterative pseudo-contrastive learning. Together, *Modules II* and *III* constitute a single training epoch, in which gene embeddings’ discriminability is enhanced. The training process continues until either the change in gene assignments falls below a threshold or a predetermined number of epochs is reached. Upon training completion, SpaCEX-generated gene embeddings (SGEs) can be utilized in downstream analytical tasks, such as i) enhancing transcriptomic coverage of FISH-based ST, ii) cross-sample gene alignment, iii) identifying disease-associated genes, iv) identifying disease-associated gene crosstalk, v) pinpointing genes with designated spatial expression patterns, vi) detecting SVGs, vii) improving spatial clustering.

### SpaCEX Identifies Spatially Co-expressed Gene Clusters as Biologically Relevant Genomic Context

The effectiveness of SpaCEX in generating semantically meaningful gene representations hinges on its ability to identify biologically relevant genomic contexts, manifested as clusters of spatially co-expressed genes. To systematically evaluate this ability, we utilize twelve human dorsolateral prefrontal cortex (DLPFC) 10x Visium datasets (10x-hDLPFC) (16) and a mouse hippocampus Slide-seqV2 dataset (ssq-mHippo) (17), as listed in *SI Appendix*, Table S1. We benchmark SpaCEX against four state-of-the-art competing methods: CNN-PReg (18), Giotto (19), Spark (20) and STUtility (21) (*SI Appendix*, Table S2). Our initial assessment involves visualizing spatial expression maps of four randomly selected genes from each of two SpaCEX-identified clusters, chosen to represent high and medium quality clusters, respectively (see *“Identifying groups of spatially co-expressed genes”* in Methods). *SI Appendix*, Fig. S1 shows that genes within both clusters exhibit congruent expression patterns. Next, the overall co-expression and spatial coherence of SpaCEX-generated clusters are quantitatively measured using two Davies-Bouldin (DB) indices (see *“Evaluation metrics”* in Methods). Fig. 2*A* shows that SpaCEX consistently outperforms the competing methods in both DB indices, demonstrating its effectiveness in identifying spatially co-expressed gene clusters.

**Fig. 2.**
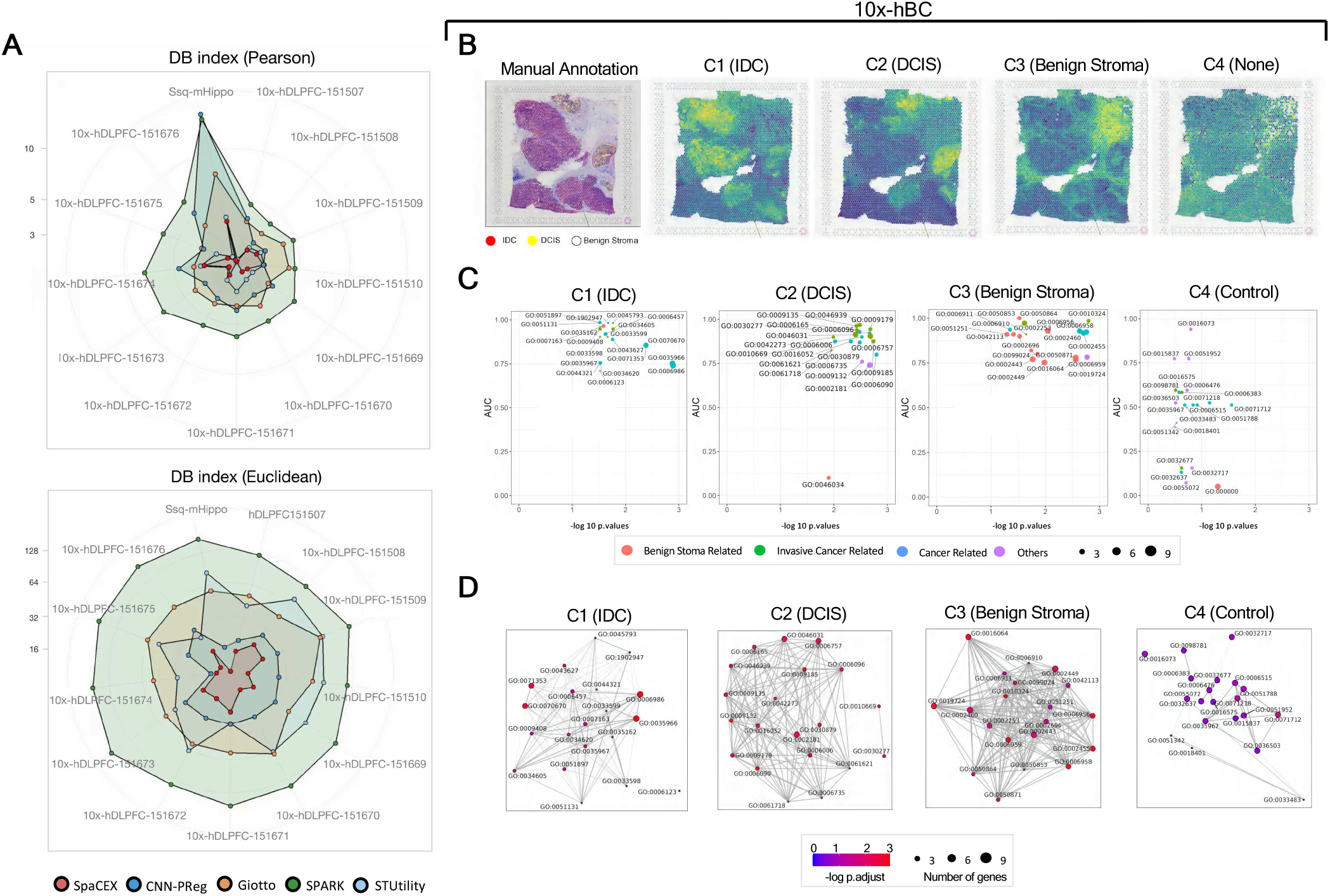
SpaCEX identifies clusters of spatially co-expressed genes within a biologically relevant genomic context. (*A*) Performance comparison of SpaCEX and four benchmark methods in grouping co-expressed genes in the 10x-hDLPFC-151673 and ssq-mHippo datasets. The DB index calculated based on Pearson and Euclidean distance are used to measure the overall co-expression (left panel) and spatial coherence (right panel) of the gene clusters, respectively. In both cases, a lower DB index value indicates enhanced co-expression or spatial coherence. (*B*) Spatial expression patterns of SpaCEX-generated gene groups overlap with the cell type distributions in the 10x-hBC dataset. The leftmost panel displays the manually annotated distributions of ductal carcinoma in situ (DCIS, in yellow), invasive ductal carcinoma (IDC, in red) and benign stroma cells (in original color) in human breast cancer tissues. The right four panels display the aggregated spatial expression (module scores) of three SpaCEX-generated gene groups (C1-C3) and a control cluster (C4) consisting of randomly selected genes. The brightness in each panel is positively correlated with the level of aggregated gene expression. The name of the cell type, whose distribution overlaps with the aggregated expression pattern of the gene group, are indicated above each panel. (*C*) GO enrichment and cofunction analyses of genes within C1-C4. The dots represent the 20 most significantly enriched GOBPs in each gene group. The x-axis represents the negative logarithm of adjusted P-values of biological process enrichment significance, while the y-axis represents the ROC AUC scores of the 20 GOBPs in the gene cofunction analysis. Red color indicates benign stroma-related GOBPs, green color the noninvasive cancer-related GOBPs, cyan color invasive cancer-related GOBPs, and purple color other GOBPs. (*D*) The connectivity between the nodes corresponding to most significantly enriched GOBP indicates their functional associations. The node color represents the GOBP’s enrichment significance (negative logarithm of adjusted P-value), with darker colors indicative of lower significance levels. Node size indicates the number of genes involved in the GOBP.

To verify the legitimacy of using SpaCEX-identified gene clusters as spatial genomic contexts, we delve into their biological significances through gene pathway enrichment analysis and gene cofunction analysis using the 10x-hBC dataset derived from human breast cancer tissue (20). Our analysis targets three specific gene clusters (C1, C2, and C3), each comprising over 20 genes and demonstrating the lowest group closeness centrality (*SI Appendix*, Supporting Text). This centrality metric suggests their expression patterns are most likely to diverge from the majority, probably due to pathological functions. As depicted in Fig. 2*B*, the aggregated expression patterns (22) (*SI Appendix*, Supporting Text) of C1 through C3 are associated with the spatial distributions of invasive ductal carcinoma (IDC), ductal carcinoma in situ (DCIS) and benign stroma cells, respectively. In contrast, the aggregated expression of C4, a control cluster comprising 30 randomly selected genes, is dispersed over the spatial map. Additionally, Fig. 2*C*-*D* showcase that these clusters are statistically significantly enriched with pathologically/biologically relevant and densely inter-connected gene ontology biological processes (GOBP). The functional coherence of member genes within these clusters is also notable, given the involvement of a gene in a GOBP can be reliably predicated based on other member genes’ involvement in the same GOBP (Fig. 2*C*). A more detailed explanation of the methodology and results of this analysis is available in the *SI Appendix*, Supporting Text. These findings altogether affirm that SpaCEX-identified gene clusters are cofunctional and biologically/pathologically relevant to the context under investigation, thus endorsing their roles as spatial genomic contexts.

### SGEs Effectively Capture Fundamental Gene Semantics

We begin this section by validating the functional and relational semantics of SGEs through three analyses (see *“Evaluating SpaCEX-generated gene embeddings”* in Methods). In the first analysis, hierarchical clustering is performed on SGEs for the KRT-II, HLA-I and HLA-II gene family members derived from human DLPFC tissues (Fig. 3*A*). We find that SGEs from the same family are clustered together, and those from functionally related gene families (e.g., HLA-I and HLA-II) are positioned in closer proximity within the hierarchy than those from less related families (e.g., KRT-II and HLA-II). In contrast, these gene families are more intermingled when the clustering is based on their original expression profiles (*SI Appendix*, Fig S2). In the second analysis, Leiden is utilized to identify gene clusters at various resolutions, using either SGEs or the original gene expression profiles. Subsequently, pathway enrichment analysis against the Reactome database (23) is conducted on these gene clusters to compile their high-confidence “pathway hits”, which is indicative of the gene representations’ efficacy in encoding complex gene-gene connections (9). Fig 3*B* showcases that gene clusters derived from SGEs yield a consistently higher average number of “pathway hits” compared to those derived from original gene expression profiles across resolution levels. The third analysis focuses on predicting gene-gene interactions using four types of gene embeddings, including SGEs derived from the 10x-hDLPFC-151676, scBERT embeddings, Gene2vec embeddings, and randomly generated embeddings. As elaborated in *SI Appendix*, Supporting Text, SGE-based prediction proves to be the most accurate, as evidenced by its superior accuracy and AUC scores (Fig 3*C*), along with a prediction heatmap close aligning with the ground truth (Fig 3*D*). Collectively, these analyses affirm that fundamental functional and relational semantics of genes are encapsulated into SGEs.

**Fig. 3.**
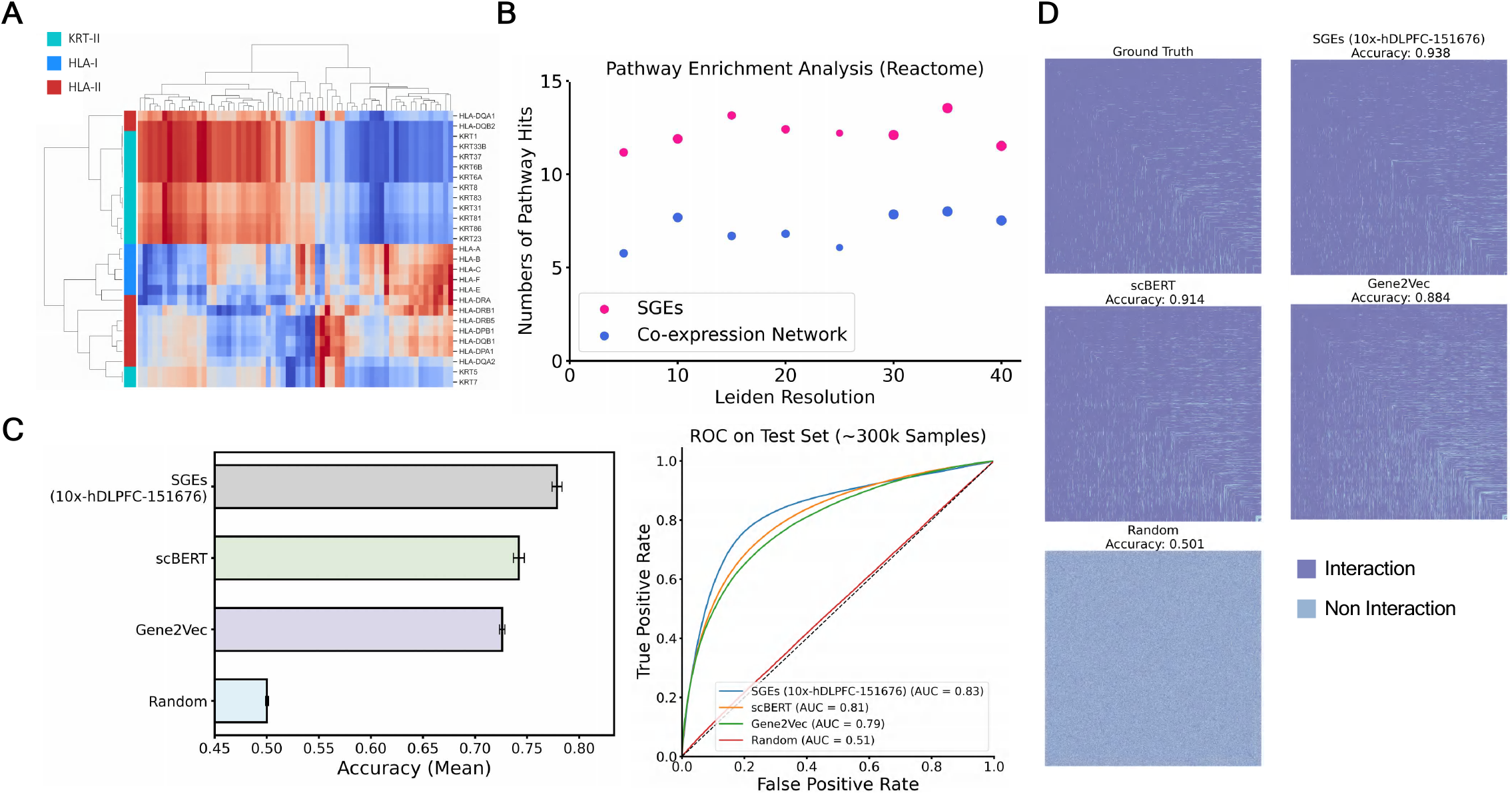
Validating the relational semantics of SGEs. SGEs are derived from the 10x-hDLPFC-151676 dataset. (*A*) Hierarchical clustering based on the SGEs of the HLA-I, HLA-II, and KRT-II gene family members. Genes with similar SGEs are positioned in close proximity along the y-axis. The 64 dimensions of the SGEs are represented along the x-axis. Gene families are indicated in different colors on the y-axis. The red color intensity in the diagram positively correlates with the SGE values. (*B*) Reactome-based pathway enrichment analysis of gene clusters generated by Leiden at various resolutions, based on gene-gene similarity matrices computed from either the SGEs or original gene expression profiles. The x-axis denotes the Leiden resolution, while the y-axis represents the average number of statistically significant enriched gene pathways (or high-confidence “pathway hits”) across the gene clusters. Red and blue spots represent gene clusters derived from the SGEs and original gene expression profiles, respectively. The predictive power of gene-gene interactions with different types of gene embeddings. Here, we showcase the mean accuracies (left panel) and ROC AUC (right panel) scores for Gene-Gene Interaction Predictor Neural Network (GGIPNN)-based predictions of gene-gene interactions using four distinct types of gene embeddings: SGEs, scBERT embeddings, Gene2vec embeddings, and randomly generated embeddings. Refer to Supplementary Note 1.4 for details of creating training and testing datasets. The gene-gene interaction heatmaps are presented to compare the ground truth (top-left) with prediction results using the four types of gene embeddings. For better visualization, the heatmaps only include the top 1,000 genes that exhibit the most interactions with other genes as per ground truth. In these maps, a filled cell in the indicates the existence of an interaction between the pair of genes in the corresponding row and column, while a blank cell indicates the opposite. The prediction accuracy is indicated on top of each heatmap.

### SGEs Facilitate Cross-sample Gene Alignment

Given that gene-gene relationships remain relatively stable across datasets under identical conditions and are less prone to technical artifacts like batch effects (24), we posit that SGEs, generated based on gene relational semantics as previously validated, are also resilient to technical artifacts, thus capable of facilitating cross-sample gene alignment. To verify this point, SGEs from two healthy human middle temporal gyrus (MTG) 10x Visium datasets (10x-hMTG-1-1 and 10x-hMTG-18-64) are aligned using our SGE alignment network (SAN), which is a three-layer feedforward neural network (FFN) with nonlinear activation functions (Fig. 4*A*). SAN learns a mapping function, F, that minimizes the mean absolute errors (MAE) between SGE pairs of 2177 housekeeping genes from two different datasets (see *“Data and Code Availability”* for where housekeeping genes are acquired). The consistent biological roles of these house-keeping genes suggest that their SGEs should align accurately if they are unaffected by cross-sample technical variations. For comparison, gene expressions from both datasets are also transformed into uniform manifold approximation and projection (UMAP) embeddings with the same dimensionality as SGEs. SAN is then trained to align these UMAP embeddings, which could potentially be confounded by technical variations. The cosine dissimilarity of aligned embeddings pairs serves as an indicator of alignment discrepancies. Fig. 4*B* reveals that discrepancies between SGE pairs are significantly lower than those between pairs of UMAP embeddings, highlighting SGEs’ robustness against cross-sample technical noises. Moreover, by visualizing aligned SGEs and UMAP embeddings on a principal component analysis (PCA) plot (Fig. 4*C*), we find SGEs from different datasets are more evenly mixed compared to the UMAP embeddings. These observations provide strong evidence to that SGEs prioritize capturing genuine gene semantics and are resistant to technical artifacts across samples, thereby facilitating more accurate cross-sample gene alignment.

**Fig. 4.**
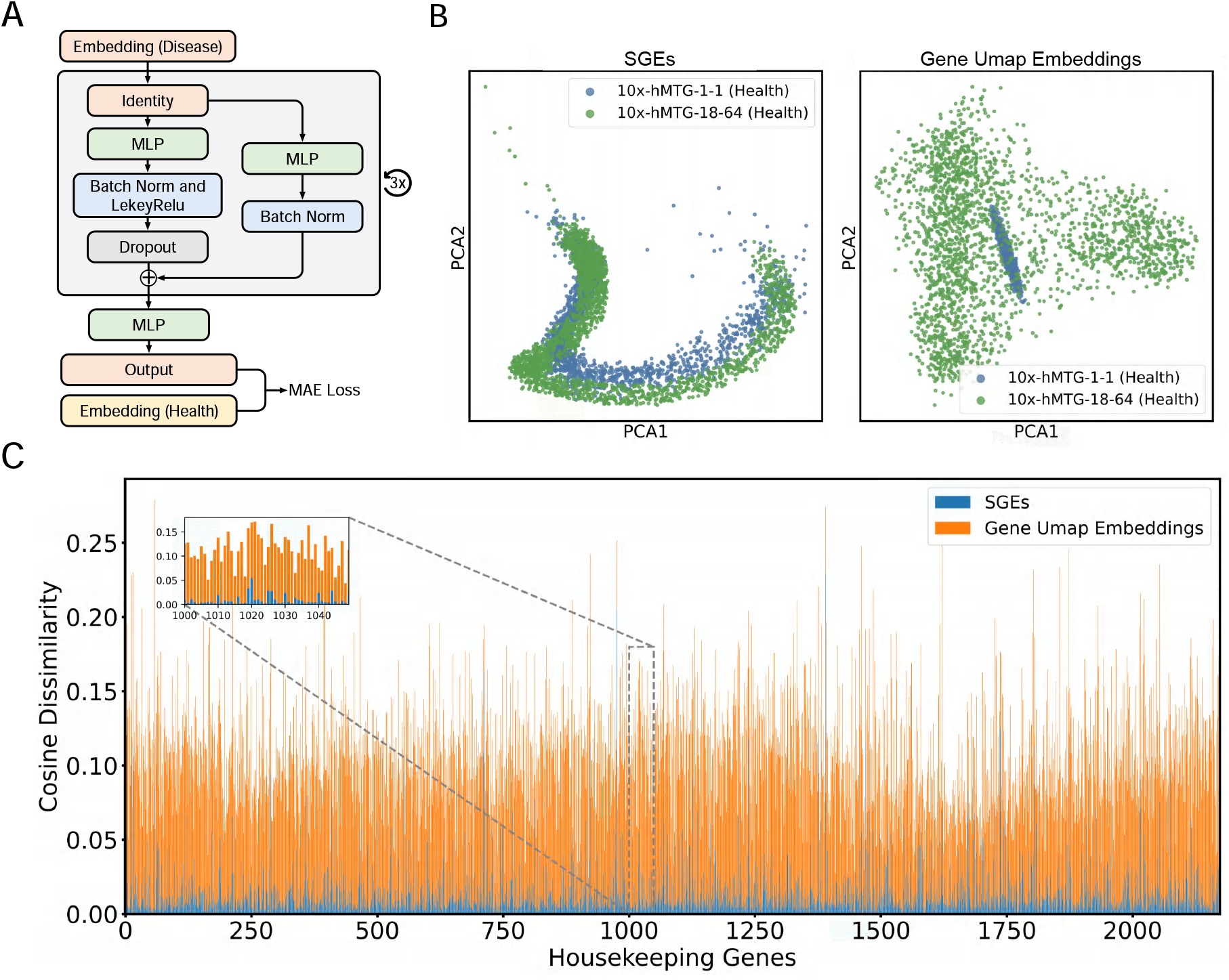
SGEs of 2177 housekeeping genes are generated from two healthy human MTG 10x Visium dataset (10x-hMTG-1-1 and 10x-hMTG-18-64), respectively. UMAP embeddings of the same dimension as SGEs for these housekeeping genes are also generated from both datasets. Embedding pairs of identical genes but from different datasets are then subjected to alignment. (*A*) The network architecture of SGE alignment network (SAN). (*B*) PCA plots of SGEs (left) and gene UMAP embeddings (right) after SAN-mediated alignment. (*C*) Scaled cosine dissimilarities between pairs of aligned SGEs (in blue) versus those between aligned UMAP embeddings (in orange).

### Identifying Disease-associated Genes and Gene Crosstalk with SGEs

Identifying genes with altered spatial expression and interactions is pivotal for illuminating pathogenic mechanisms underlying disease progression, e.g., the elevated APOE expression within hippocampus in Alzheimer’s Disease (AD) (25) and the intensified interplay between Notch and Wnt pathways in many cancers (26). We define strategies for identifying disease-related genes as either reference-based or reference-free. The former compares spatial gene expressions in diseased versus healthy tissues to pinpoint differences, while the latter identifies genes exhibiting specific expression patterns within putative pathogenic regions independently of healthy tissue expression benchmarks. Herein, we detail the use of SpaCEX and SGEs in both strategies.

#### Reference-based

In ST, directly comparing gene expression patterns between conditions is difficult due to variations in tissue slice preparations, technical artifacts, and spatial heterogeneity across slices. Nevertheless, SGEs, which are context-aware embeddings capturing fundamental gene semantics and whose value discrepancies across samples/conditions are reconcilable by SAN-mediated gene alignment as shown in Fig. 4*A*, allow for the detection of disease-associated alterations in spatial gene expressions and relationships.

To verify this, we generate SGEs from two healthy brain tissue datasets (10x-hMTG-1-1 and 10x-hMTG-18-64) and one AD brain tissue dataset (10x-hMTG-2-3). SAN described in the preceding section is used to align SGE pairs of identical genes from two different datasets. The 2177 housekeeping genes are randomly split into an “anchor” set of 2051 genes for SAN training and a “non-anchor” set of 126 genes reserved for testing. The trained SAN is applied to align SGE pairs from both the non-anchor housekeeping gene set and a set of 42 AD-associated genes reported by previous studies (*SI Appendix*, Table S3). When aligning SGEs between an AD and a healthy dataset (i.e., 10x-hMTG-1-1), we find that AD-associated genes demonstrate significantly greater PCA distances (Fig. 5*A* *Top*) and scaled cosine dissimilarities (Fig. 5*B* *Top*) compared to the non-anchor housekeeping genes. In contrast, alignment of SGES between the two healthy datasets does not show such disparities, neither in PCA distances (Fig. 5*A* *Bottom*) nor in scaled cosine dissimilarity (Fig. 5*B* *Bottom*), aligning with expectations that biological semantics of AD-related genes should remain unchanged in healthy conditions. These findings imply that differences in SGEs between healthy and diseased states could signal changes in gene functions associated with the disease. Therefore, genes can be prioritized based on the degree of their SGE dissimilarities, furnishing insights for the identification of disease-associated genes.

**Fig. 5.**
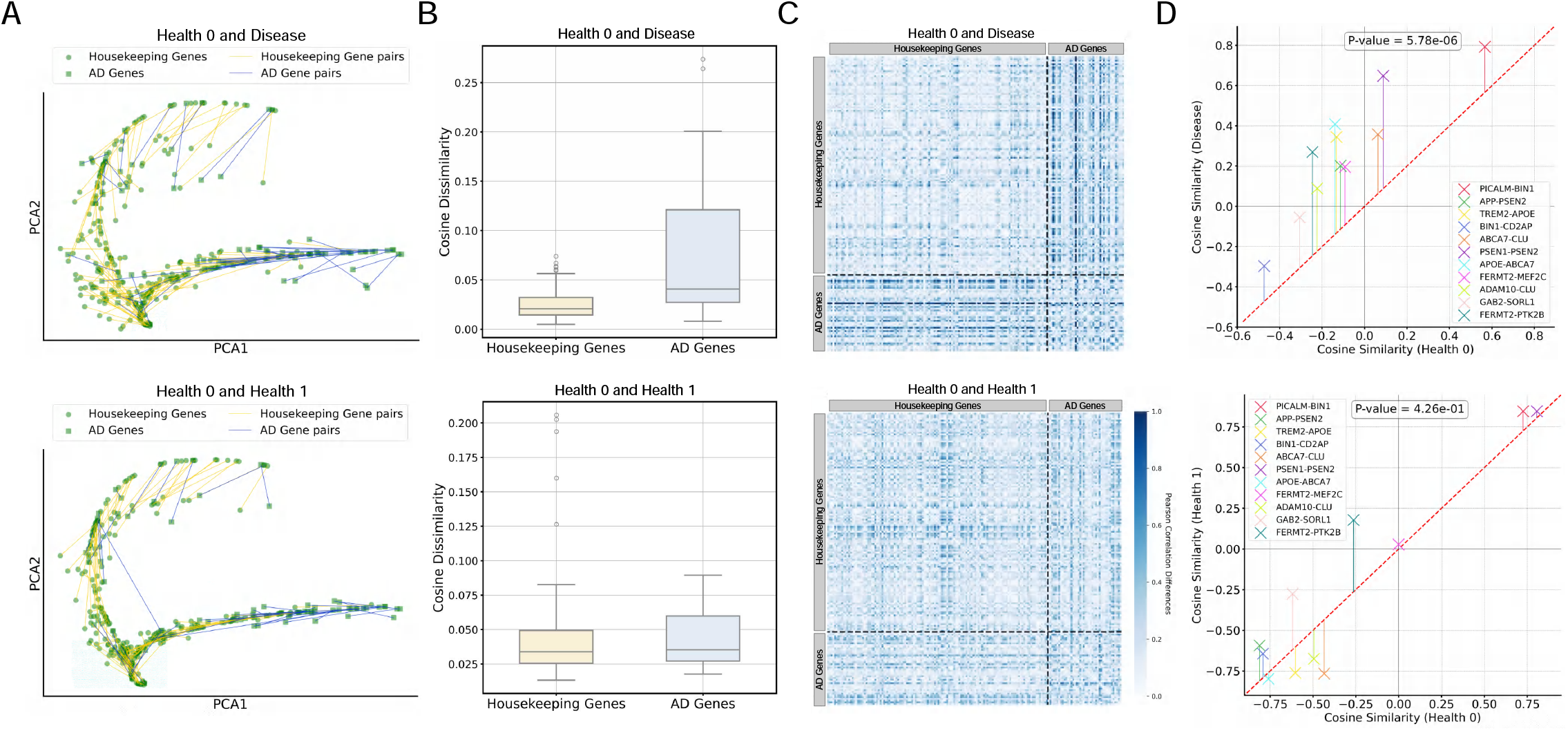
Identifying AD-associated genes and gene crosstalk with SGEs. Fig. 5*A* and 5*B* are for identifying AD-associated genes, while Fig. 5*C* and Fig. 5*D* for AD-associated gene crosstalk. SGEs of 42 AD-associated genes and 126 non-anchor housekeeping genes are obtained from two healthy MTG 10x Visium datasets (health_0: 10x-hMTG-1-1 and health_1: 10x-hMTG-18-64) alongside an AD MTG 10x Visium dataset (10x-hMTG-2-3). SGE pairs of identical genes but from different datasets are aligned using SAN. The AD and the healthy (health_0) datasets form the study group, while the two healthy datasets form the control group. (*A*) PCA plots of SGEs in the study (left) and control (right) groups. Round dots and triangles represent housekeeping and AD-associated genes, respectively. The PCA distances between SGE pairs are visually represented by yellow and blue lines for housekeeping and AD-associated genes, respectively. Compared to the control group, blue lines are markedly longer than yellow lines in the study group. (*B*) Box-plots of scaled cosine dissimilarities between SGE pairs in the study (left) and control (right) groups. Yellow and blue boxes represent housekeeping genes and AD-associated genes, respectively. (*C*) Gene-gene interactions within each dataset are quantified using a Pearson correlation matrix calculated from SGEs. Alterations in gene-gene interactions between two datasets are measured as a correlation shift matrix representing the absolute differences between the two correlation matrices, which is visualized as a heatmap wherein darker colors indicates larger shifts. Compared to the control group (right), correlation shifts in the study group (left) for gene pairs involving at least one AD-associated gene are significantly larger than those for gene pairs of housekeeping genes only. (*D*) Scatterplots of correlations between gene pairs involved in the same AD-associated pathways, with each cross representing a gene pair. For the study group (left), y- and x-axes denote gene correlations in the AD and healthy (health_0) datasets, respectively. In the control group (right), these axes represent gene correlations in the health_1 and health_0 datasets, respectively. A t-test is used to assess the statistical significance of differences in gene correlations between the datasets, with P-values indicated on top of each panel.

SGEs can also provide valuable clues to altered gene crosstalk in disease. We denote the Pearson correlations between SGEs of gene pairs from a healthy brain tissue dataset (10x-hMTG-2-5) as *ρ*_*health*_, and from an Alzheimer’s Disease (AD) dataset (10x-hMTG-2-3) as *ρ*_*AD*_. Fig. 5*C* reveals that the absolute difference between these correlations, *δ*_*ρ*_=|*ρ*_*health*_ *− ρ*_*AD*_|, signals potential alterations in gene crosstalk, as *δ*_*ρ*_ values are notably higher when at least one gene in the pair is AD-associated compared to pairs of house-keeping genes. This observation is bolstered by a statistically significant increase (P-value=5.78e-06) in correlations between gene pairs that participate in the same AD-related pathways (Fig. 5*D* *Top*) in the AD dataset. In contrast, no such shifts in correlation are detected between the two healthy datasets (Fig. 5*D* *Bottom*), in line with the expectation that gene relationships in health conditions should remain largely stable. Therefore, substantial changes in SGE correlations are indicative of disease-related alterations in gene interactions.

#### Reference-free

Our reference-free approach involves employing our innovative method, SpaCEX-SPS, to create a pseudo-gene with designated spatial expression patterns (see *“SpaCEX-SPS”* in Methods), subsequently converted into an SGE. Based on the scaled cosine similarities between the SGEs of real genes and the pseudo-gene, we can pinpoint genes whose spatial expression patterns correspond to the predefined one. Our method is tested on two datasets: a human breast cancer dataset (10x-hBC) and an AD MTG dataset (10x-hMTG-2-3). In the SpaCEX-SPS simulation, “high”-level of gene expression is defined as the 95th percentile of average expressions across all genes, “medium”-level the 75th percentile, and “low”-level the 35th percentile. For the 10x-hBC dataset, we aim to discover genes with high expression within tumor cores (IDC), medium expression within tumor edges, and low expression elsewhere. Using SpaCEX-SPS, we simulate a gene mirroring these expression patterns. As shown in Fig. 6*A*, we successfully identify cancer-associated genes such as BRCA1, BRCA2, PALB2, and TCEAL4, all exhibiting the sought-after expression patterns. Similarly, for the 10x-hMTG-2-3 dataset, we pinpoint AD-associated genes like TREM2, PSEN1, BIN1, and APOE, characterized by high expression within the white matter (WM) layer and medium expression within other cortex layers (Fig. 6*B*). Particularly, the upregulation of BIN1 within the WM of AD brain has been previously reported (27). These outcomes collectively demonstrate the efficacy of utilizing SGEs and SpaCEX-SPS for discerning genes with disease-specific spatial expression profiles. Moreover, our methodology extends beyond disease-associated gene identification to encompass any genes with designated spatial expression patterns.

**Fig. 6.**
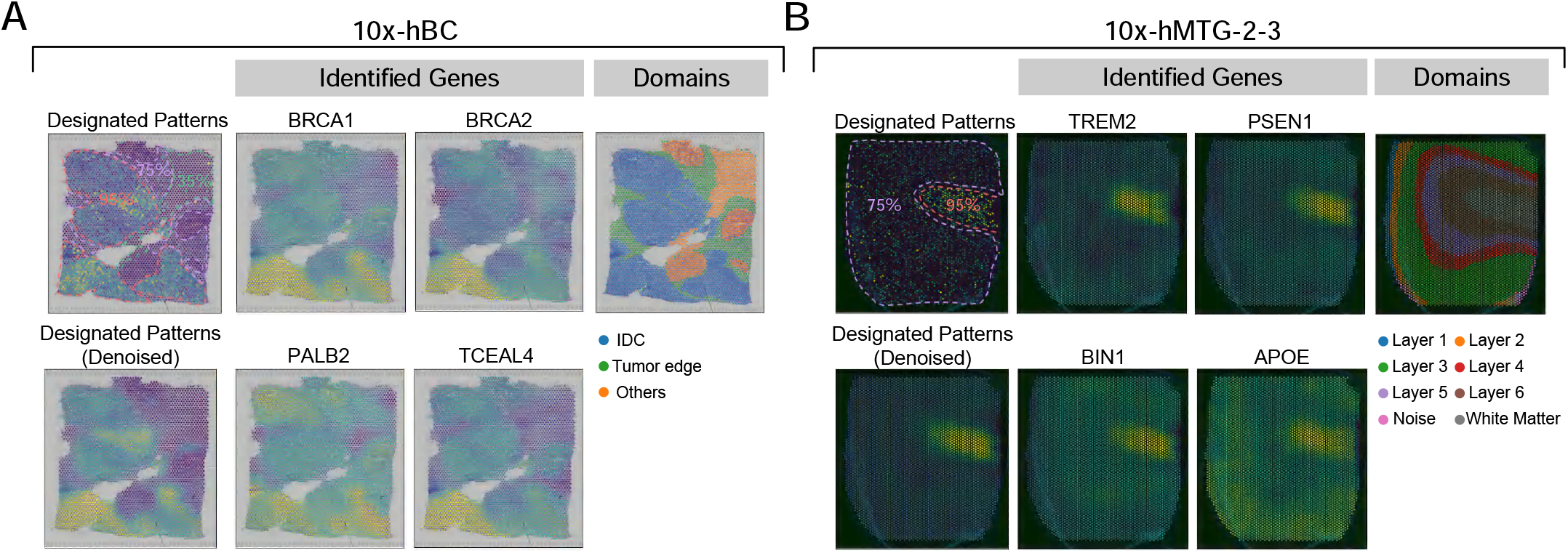
Identifying disease-associated genes with designated spatial expression patterns. For both Fig. 6*A,B*, the designated gene spatial expression patterns are displayed in the leftmost panel in the first row. The expression level percentiles are indicated within different regions demarcated by dotted lines of varying colors. The rightmost panel in the first row show expert-curated domain labels. The leftmost panel in the second row represents denoised SpaCEX-SPS-simulated genes that mirror the designated patterns. The rest panels correspond to denoised spatial maps of identified real genes whose expression patterns resemble the designated ones. (*A*) The designated expression patterns have a high expression level (95% percentile) in tumor cores (i.e., IDC), a medium expression (75%) in tumor edges, and a low expression (35%) elsewhere in the human breast cancer 10x Visium dataset (10x-hBC). (*B*) The designated expression patterns have a high expression level (95% percentile) in white matter (WM), a medium expression (75%) in other cortex layers in the human MTG 10x Visium dataset (10x-hMTG-2-3).

### SGE-based Enhancement of the Transcriptomic Coverage in FISH-based ST

A significant challenge in ST is achieving both full transcriptomic coverage and high-resolution. Existing methods like Tangram (28), SpaGE (29) SpaOTsc (30) augment transcriptomic coverage in high-resolution, FISH-based ST data, but they essentially generate “pseudo-ST” data since they focus on mapping single cells in scRNA-seq onto spatial locations in ST to compensate for genes not profiled in ST with scRNA-seq data, rather than the *de novo* generation of ST data with inherent spatial semantics. These methods are limited by their underutilization of spatial information in the mapping process, as seen in Tangram and SpaGE, and the introduction of systematic biases from discrepancies between scRNA-seq and ST data such as inconsistencies in data scales.

To address this challenge, we introduce SpaCEX-enhanced-transcriptomics-coverage (SpaCEX-ETC), an innovative SGE-based Generative Adversarial Network (GAN) model as detailed in the *“SpaCEX-ETC”* in Methods and Fig. 7*A*. SpaCEX-ETC is predicated on the notion that gene relational semantics should remain largely consistent across different ST data types for the same tissue type. Consequently, the spatial expression profiles of uncovered genes can be extrapolated from those of covered genes, drawing on their semantic relationships inherent in SGEs derived from a full transcriptomic coverage ST dataset (e.g., 10x Visium). We evaluate the effectiveness of SpaCEX-ETC by reproducing the 330 covered genes from a mouse embryo SeqFISH dataset (sqf-mEmb), guided by the SGEs from a mouse embryo 10x Visium dataset (10x-mEmb).

**Fig. 7.**
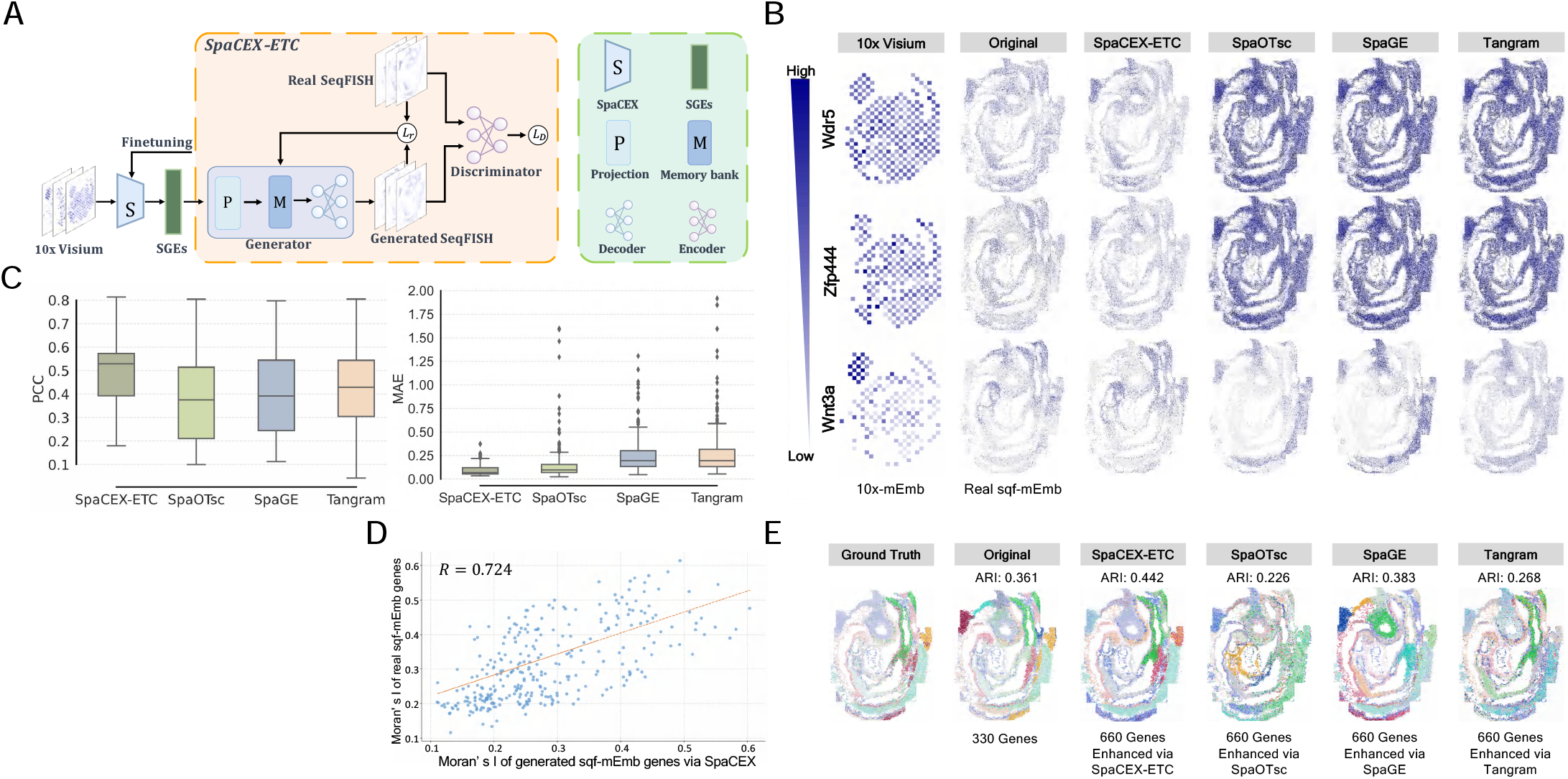
SGE-based enhancement of transcriptomic coverage of FISH-based ST with SpaCEX-ETC. (*A*) Workflow of SpaCEX-ETC. Initially, genes from a ST dataset with full transcriptomic coverage are encoded into SGEs. In the training phase, SGEs of the genes present in the target FISH-based ST dataset are fed into SpaCEX-ETC’s generator to regenerate their original spatial gene expressions. The generator consists of three components: a multilayer perceptron (MLP)-based encoder, an MLP-based decoder, and a memory bank that ensures the training stability. Following this, a discriminator is trained to distinguish between the original and regenerated genes. Meanwhile, the loss gradients from the generator are backpropagated to update the parameters of the MAE’s encoder within SpaCEX, so that SGEs are more adapted to the specific semantics inherent in the FISH-based ST dataset. Once trained, SpaCEX-ETC can generate genes that are initially absent in the dataset, using their finetuned SGEs. (*B*) Three genes (Wdr5, Zfp444, and Wnt3a), which are covered by both the SeqFish (sqf-mEmb) and the 10x-mEmb datasets and differ in their expression levels, are used to evaluate SpaCEX-ETC. These genes’ SGEs are initially obtained from the 10x-mEmb dataset and used to regenerate their expression profiles in the sqf-mEmb dataset. From the left to the right, we show the three genes’ original spatial expression profiles in the 10x-mEmb dataset, in the sqf-mEmb dataset, alongside their regenerated expression profiles by SpaCEX-ETC and the three benchmark methods. (*C*) Displayed in the box plot are the Pearson correlation coefficients (PCCs) between the original and spatial expression profiles of the 330 genes regenerated by SpaCEX-ETC and the benchmark methods. (*D*) The scatterplot shows spatial variabilities of the 330 genes and their respective regenerated counterparts, with spatial variability quantified using Moran’s I index. The red line in the scatterplot represents a fitted regression line with R=0.724. (*E*) The transcriptomic coverage of sqf-mEmb is doubled by using SpaCEX-ETC and the benchmark methods to generate an additional 330 genes absent in the original dataset. The leftmost spatial map shows the ground truth tissue domain annotations, the second map shows the spatial clustering results of SpaGCN using the unexpanded gene set, while subsequent maps showcase the spatial clustering results using gene set augmented by SpaCEX-ETC and the benchmark methods, respectively. The clustering accuracy (i.e., ARI) is shown below each method name.

We include Tangram, SpaGE, and SpaOTsc as benchmarks (*SI Appendix*, Table S2) to reproduce the same gene set from scRNA-seq data. To ease a direct comparison between the original and reproduced genes, we visualize the spatial maps of three genes of different expression levels, including Wdr5, Zfp444, and Wnt3a. Fig. 7*B* illustrates that SpaCEX-ETC surpasses benchmark methods in accurately reproducing genes, achieving high fidelity in both spatial expression patterns and data scales.

Additionally, Fig. 7*C* quantitatively demonstrates SpaCEX-ETC’s superiority in generating genes that closely correlate with their actual values. This concordance is further supported by the highly correlated spatial variability (R=0.724) between authentic and SpaCEX-ETC-generated gene expressions, as depicted in Fig. 7*D*. Finally, we select an additional 330 genes imputed by SpaCEX-ETC, deemed most real-like by the GAN discriminator, to double the transcriptomic coverage of the sqf-mEmb dataset. This augmented gene set undergoes spatial clustering to evaluate the imputed genes’ quality and their analytical utility. Fig. 7*E* shows that spatial clustering with SpaCEX-ETC-imputed genes achieves a significantly higher accuracy compared to either the original dataset or genes imputed by the benchmark methods. Collectively, these results highlight the efficacy of SGEs in enhancing the transcriptomic coverage of FISH-based ST via a generative approach.

### SGE-based SVG Detection

In this section, we introduce a novel computational method, SpaCEX-SVG, that leverages SGEs to detect SVGs from ST datasets, as detailed in the *“SpaCEX-SVG”* section in Methods. Essentially, SpaCEX-SVG calculate a spatial variability score for each gene based on the similarity of its SGE with those of simulated spatially homogeneous genes. SVGs then are ranked and selected according to these scores. For this assessment, we select the top 3000 SVGs from both the 10x-hDLPFC-151507 and 10x-hBC datasets using SpaCEX-SVG and two benchmark methods: SpatialDE and SPARK-X (31) (*SI Appendix*, Table S2). Both Moran’s I and Geary’s C indices shows that the SVGs selected from the 10x-hDLPFC-151507 dataset by SpaCEX-SVG are more spatially variable than those selected by the benchmark methods (*SI Appendix*, Fig. S3A). Moreover, one well-documented drawback of the benchmark methods is their inability to effectively rank SVGs based on their spatial variability scores (P- or Q-values) (32). In contrast, SpaCEX-SVG is more sensitive and effective in distinguishing levels of spatial variability among SVGs. This is exemplified in *SI Appendix*, Fig. S3B, where the top four SVGs selected by SpaCEX-SVG exhibit more noticeable spatial variabilities than those selected by the benchmark methods. A parallel analysis conducted on the 10x-hBC dataset yields similar results (*SI Appendix*, Fig. S4). These results altogether demonstrate the potential of SGEs for detecting and ranking SVGs in ST datasets.

### SGE-improved Spatial Clustering

In this section, we propose SpaCEX-Improved-Spatial-Clustering (SpaCEX-ISC), a novel SGE-based computational method for enhancing spatial clustering, as detailed in the *“SpaCEX-ISC”* section in Methods. The rationale behind SpaCEX-ISC is that spatial clustering can be effectively improved by optimizing the informational efficiency of spatial transcriptomic data, which involves minimizing redundant information and retaining the most discriminative information presented by feature genes (33).

Redundant information among genes can be revealed through their SGE similarity matrix and reduced by only selecting and retaining the most discriminative genes within groups of highly similar ones. The most discriminative genes are those with the highest spatial variability scores, as determined by SpaCEX-SVG. Subsequently, any spatial clustering algorithm, which is SpaGCN in our case, can work with this information-efficient set of feature genes to achieve improved performance. We select two state-of-the-art spatial clustering methods, GraphST and SpaGCN, alongside a baseline method, Leiden, as benchmarks for comparison (*SI Appendix*, Table S2). In a comprehensive evaluation across twelve 10x-hDLPFC datasets, SpaCEX-ISC consistently outperforms the benchmark methods, as evidenced by its highest Adjusted Rand Index (ARI) and Normalized Mutual Information (NMI) scores (*SI Appendix*, Fig. S5A). SpaCEX-ISC’s superiority over the benchmark methods is further illustrated by its more accurately recovered annotated anatomical cortex layers in the spatial maps of the 10x-hDLPFC-151676 and 10x-hDLPFC-151669 datasets (*SI Appendix*, Fig. S5B). Finally, SpaCEX-ISC achieves optimal performances across six 10x-hDLPFC datasets when approximately 50%-60% redundant information is excluded (*SI Appendix*, Fig. S5C).

## Discussion

In ST, Genomic contexts unveiled as groups of spatially cofunctional and co-expressed genes are instrumental for generating semantically rich gene embeddings, paralleling the concept of word vectorization in natural languages processing. Existing foundational models designed to learn gene embeddings from microarray or scRNA-seq data typically rely on massive pretraining, resulting in a weakened sensitivity to context-specific nuances, and fall short of incorporating spatial expression information into the gene vectorization process. In this work, we propose SpaCEX, a novel context-aware, self-supervised learning model that exploits spatial genomic contexts in ST to derive distributed gene representations imbued with spatial gene functional and relational semantics.

We comprehensively evaluate SpaCEX across ST datasets of various tissues, species, and platforms in aspects regarding the model’s legitimacy and its utility in downstream, task-specific applications. To establish the methodological soundness, we initially demonstrate SpaCEX’s adeptness at identifying spatial genomic contexts as groups of cofunctional and co-expressed genes. Subsequent analyses confirm SpaCEX’s ability in generating SGEs that encapsulate essential gene semantics from these genomic contexts, with SGE correlations reflecting gene familial and ontological ties. Notably, SGEs prioritize biological variations over technical noises, enhancing their utility in cross-sample gene alignment, as demonstrated by the accurate alignment of SGEs of functionally stable house-keeping genes. For task-specific applications, we propose a suite of innovative SGE-based methods for identifying disease-associated genes and gene crosstalk, pinpointing genes with designated spatial expression patterns, enhancing the transcriptomic coverage of FISH-based ST, detecting SVGs, and improving spatial clustering. These methods either pioneer solutions to existing problems or markedly surpass established benchmarks.

SpaCEX’s remarkable performance are rooted in four aspects: the effective integration of spatial expression patterns into SGEs via an image-focused, self-supervised MIM approach; learning relational semantic structures among genes through a flexible and robust SMM-based clustering; a novel combination of MIM with contrastive learning for iterative joint optimization, enhancing the perceptibility and discriminability of SGEs; and the resilience of SGEs to technical noise, ensuring reliable gene alignment across conditions. Overall, SpaCEX not only facilitates the discovery of cofunctional gene modules, like gene networks and pathways, but also generates biologically significant gene embeddings, laying the foundation for a suite of downstream task-specific tools. Thus, SpaCEX promises to contribute to the development of a genomic “language”-based methodological ecosystem. Future improvements for SpaCEX may include enriching the informativeness of SGEs through a multimodal learning approach, integrating diverse gene relational semantics from additional datasets like gene co-expression patterns across cell types observed in scRNA-seq.

## Methods

### Data Quality Control and Preprocessing

We conform to the conventional procedure for preprocessing ST data, as implemented in the S CANPY package (34). Specifically, we first remove mitochondrial and External RNA Controls Consortium (ERCC) spike-in genes. Then, genes detected in fewer than 10 spots are excluded. To preserve the spatial data integrity, we do not perform quality control on spatial spots. Finally, the gene expression counts are normalized by library size, followed by log-transformation.

### Representation Learning of Spatial Gene Expression Maps

As the spatial gene maps can be visualized as gray-scale images, we devise an adapted version of MAE to transform visual features of gene images into embeddings in a latent feature space. A given gene image is first segmented into regular non-overlapping patches, from which a subset of patches is randomly selected, masked and discarded. The remaining patches are fed into the MAE encoder to generate visible patch embeddings. Given that a gene image is gray-scale and often sparse, we use a higher masking ratio (80%) and a light-weighted ViT encoder with four transformer blocks and four attention heads rather than the masking ratio (75%) and the ViT-L encoder in the original paper. The visible patch embeddings and trainable tokens of masked patches are then input into the MAE decoder to reconstruct the gene image. We replace the transformer architecture of the original MAE decoder with a convolutional autodecoder to enhance the performance in our case. A more important modification is the adding of a nonlinear projection head to the end of the encoder. This projection head consists of a linear layer, a batch normalization (BN) layer and a Scaled Exponential Linear Unit (SELU) activation layer. Owing to the BN layer and the self-normalizing property of the SELU function, the gene embeddings output from the encoder more closely conform to the mixed Student’s t distribution.

### SMM-based Modeling

As Stuhlsatz et al (35) have demonstrated the capability of deep image encoder in learning visual representations that follow a multivariate Student’s t-distribution, we utilize an SMM to model the distributions of SGEs in a latent feature space, with individual components of the SMM corresponding to distinct gene clusters. The key strength of this modeling is its robustness to outliers, which are assigned reduced weights during the estimation of model parameters. Specifically, let *Z ∈ R*^*N×D*^ denote SGEs, where *N* is the total number of genes and *D* is the dimension of the feature space. We model the distribution of Z as an SMM parameterized by Θ = *{*Θ_*k*_:*π*_*k*_, *µ*_*k*_, Σ_*k*_, *v*_*k*_, *∀k ∈ K}*. Here, K represents the total number of gene clusters, while *π*_*k*_, *µ*_*k*_, Σ_*k*_, *υ* _*k*_ represents the weight, mean, covariance matrix and freedom of the *k*-th component, respectively. The density function of *z*_*i*_ is then formulated as follows:

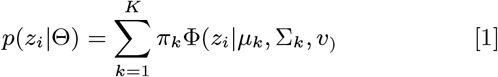

We utilize an Expectation-Maximization (EM) algorithm to iteratively estimate parameters of the SMM. Given a multitude of parameters in the model, a conventional MLE-based EM tends to overfit the data. To mitigate this problem, we introduce priors on the model parameters for model regularization purpose: we use a conjugate Dirichlet prior on Π and a normal-inverse Wishart (NIW) prior on *µ*_*k*_,Σ_*k*_:

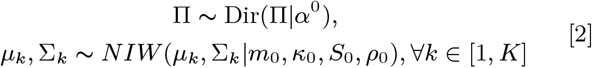

To simplify the EM algorithm, we rewrite the Student’s t distribution as a Gaussian scale mixture by introducing an “artificial”hidden variable *ζ*_*i,k*_, *∀i ∈* [1, *N*], *∀k ∈* [1, *K*] that follows a Gamma distribution parameterized by *v*_*k*_:

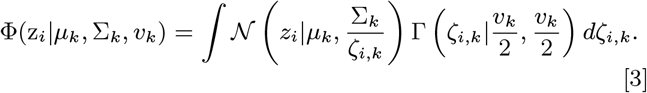

We also introduce a missing variable *ξ*_*i*_ to represent the component membership of z_*i*_. Then the posterior complete data log likelihood can be written as:

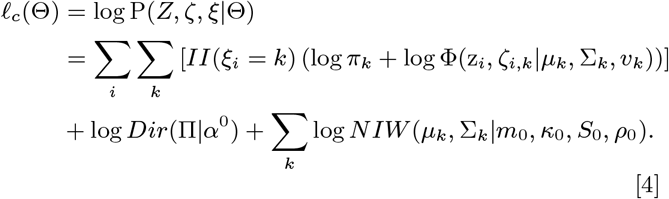

In the *t*-th iteration of the E-step, the expected sufficient statistics 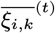 and 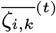 are derived based on Θ^(*t−*1)^. In the subsequent M-step, Θ^(*t−*1)^ is updated to Θ^(*t*)^ by maximizing the auxiliary function *Q*(Θ, Θ^(*t−*1)^) = *E*(*𝓁*_*c*_(Θ)|Θ^(*t−*1)^). Note that 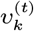 is estimated via a Generalized EM (GEM) technique to speed up the calculation without harming its converging to at least a local optimum. The two steps are alternatively conducted until either convergence is achieved or a pre-specified maximum number of iterations is reached. Refer to *SI Appendix*, Supporting Text for details about the model inference.

### Self-paced Pseudo-contrastive Optimization of SGEs

Two loss functions, *ℒ* _1_ and *ℒ* _2_, are calculated based on clustering results for updating parameters of both representation learning and the SMM through loss gradient backpropagation. This iterative process progressively improves the clustering-oriented image embeddings and clustering results. Upon completing the inference of SMM parameters 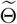 in each epoch, let *W* and 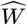 represent the parameters of the encoder and decoder of the representation learning model respectively, an epoch-level loss *ℒ* _1_ is calculated for updating parameters of MAE :

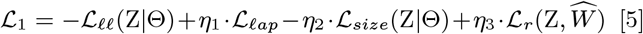

Here, *ℒ* _*𝓁ap*_ is a Laplacian regularization term that promotes the similarities among image embeddings Z to be consistent with a seeding image-image similarity matrix *𝒮*, informing the initial training phase. The derivation of *𝒮* is detailed in *SI Appendix*, Supporting Text. *ℒ* _*𝓁ap*_ is defined as follows:

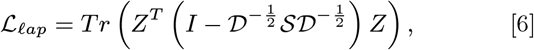

where *𝒟* is the degree matrix of *𝒮*, and *η*, initially set at 0.5, decays over the training course so that the influence of *𝒮* is gradually reduced. *ℒ* _*𝓁𝓁*_ represents the log likelihood of the embeddings given the estimated SMM parameters 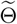 :

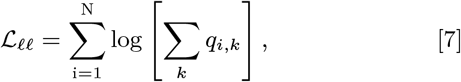

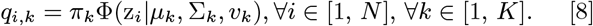

*ℒ* _*size*_ penalizes empty and tiny clusters, while exempting those whose size exceeds a predefined threshold *?* so that image assignments is not overly uniform:

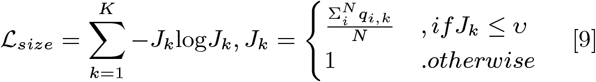

*ℒ* _*r*_, represents the fidelity loss of the reconstructed gene image by the convolutional autodecoder, expressed as:

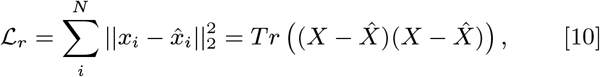

where 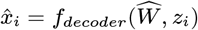 This term supervises the training of encoder, guiding it to generate gene embeddings that preserves the local structural integrity of spatial gene expressions. We set *η* _1_=0.5, *η* _2_=0.1, *η* _3_=0.1. Note, the value of *η* _1_ decays as the training progresses so that the impact of the seeding matrix diminishes over the training course.

Subsequently, within the same epoch, we utilize a batch-level loss:

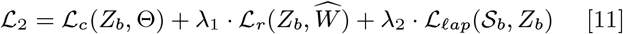

to update MAE and SMM parameters across successive batches. Here, *ℒ* _*r*_ and *ℒ* _*𝓁ap*_ remains same as in Equations Equation (10) and Equation (6) except being calculated on the batch-level. *ℒ* _*c*_ boosts high-confidence images, incrementally grouping similar instances while separating dissimilar ones:

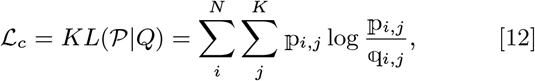

where 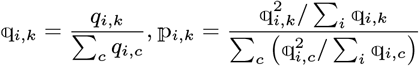

Here, *q*_*i,k*_ is same as in Equation 8, 𝕢_*i,k*_ represents the probability of assigning *i*-th gene to the *k*-th SMM component, and IP_*i,k*_ an auxiliary target distribution that boosts up high-confidence images. After this joint optimization, the training progresses to the next epoch, iterating until the end of the training process. The mathematical derivations of gradients of *ℒ* _1_ and *ℒ* _2_ with respect to *W*, 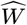 and Θ are detailed in *SI Appendix*, Supporting Text.

### SpaCEX-SPS

The section describes a generalized linear model (GLM)-based method, SpaCEX-spatial pattern simulator (SpaCEX-SPS), for simulating genes with specific spatial expression patterns in a dataset. It uses a negative binomial (NB) distribution to model gene expressions at various spots, incorporating mean and dispersion parameters that relate to the variance and squared coefficient of variation. The model estimates these parameters through regression analysis, predicting spatial expression levels at desired quantiles in specific tissue regions. By ranking and assigning expression levels based on the NB distribution, the method ensures that the simulated genes reflect the spatial structure inherent in the ST dataset, allowing for precise control over the expression pattern of simulated genes across different spatial regions. For further details, see the *SI Appendix*, Supporting Text.

### SpaCEX-ETC

As shown in Fig. 7A, SpaCEX-ETC is a GAN-based model that uses an encoder to transform gene expression matrices into SGEs, which then serve as inputs for a generator. The generator, consisting of an encoder, a decoder, and a memory bank, reconstructs gene expression profiles using an attention mechanism and a continuously updated embedding queue. The discriminator, an MLP-based network, distinguishes between actual and generated gene expressions. The model adjusts the MAE encoder weights in SpaCEX’s *Module I* through adversarial and reconstruction losses, which enables the SGEs to adapt to the particular semantics inherent in the FISH-based ST dataset. This finetuning facilitates the generation of those genes uncovered in the FISH-based dataset with optimized fidelity. For additional details, refer to the *SI Appendix*, Supporting Text.

### SpaCEX-SVG

In our study, we first developed a method that simulates spatially homogeneous genes using observed spatial transcriptomics data, applying either Negative Binomial or Zero-Inflated Negative Binomial distributions. We estimate parameters directly from the data to simulate genes and generate SGEs for both real and simulated genes. These embeddings enable us to calculate spatial variability scores by comparing real genes to their simulated counterparts using scaled cosine dissimilarity. We then rank the genes by these scores to identify those with notable spatial variations. Details provided in the *SI Appendix*, Supporting Text.

### SpaCEX-ISC

In our study, we employ the SpaCEX-ISC method to optimize the analysis of spatial transcriptomics data. This involves constructing a gene identity matrix *G* to distinguish genes across functional groups, and a similarity matrix *S* to assess gene relationships based on SGEs. We then apply Shi-Malik spectral clustering on an adjacency matrix derived from *S* to organize genes into functionally coherent groups. Using SpaCEX-SVG, we calculate spatial variability scores for each gene to identify significant spatial expression variations. These scores are used to filter out redundant data, ensuring that only the most informative gene expressions are retained. The processed data is then fed into a graph neural network to generate spot embeddings for clustering, enhancing the clarity and utility of spatial gene expression analysis. For additional details, refer to the *SI Appendix*, Supporting Text.

### Experimental Settings

Detailed experimental settings for the SpaCEX study are extensively documented in the *SI Appendix*, Supporting Text. These include the methodologies for identifying groups of spatially co-expressed genes, protocols for enrichment analysis, techniques for cofunction analysis of intra-cluster genes, criteria for evaluating SpaCEX-generated gene embeddings, and the metrics used to assess the overall performance of SpaCEX.

## Supporting information

Supporting text, Figures S1 to S5, Tables S1 to S6, SI References

## Data and Code Availability

All data are available in the main text or the supplementary materials: The mouse hippocampus dataset (ssq-mHippo) can be downloaded from https://singlecell.broadinstitute.org/single_cell/study/SCP815/sensitive-spatial-genome-wide-expression-profiling-at-cellular-resolutionstudy-summary. The human dorsolateral prefrontal cortex datasets (10x-hDLPFC) are available through the spatialLIBD package (36) at http://spatial.libd.org/spatialLIBD. The human breast cancer dataset (10x-hBC) can be obtained from https://support.10xgenomics.com/spatial-gene-expression/datasets/1.1.0/V1_Breast_Cancer_Block_A_Section_1. The three human MTG datasets, including two healthy datasets (10x-hMTG-1-1 and 10x-hMTG-18-64) and an AD dataset (10x-hMTG-2-3), are available in the GEO database (GSE220442) (37). The mouse embryo dataset based on 10x Visium (10x-mEmb) can be found at https://www.ncbi.nlm.nih.gov/geo/query/acc.cgi?acc=GSE178636. The mouse embryo dataset based on SeqFISH (sqf-mEmb) is obtainable at https://crukci.shinyapps.io/SpatialMouseAtlas/. The detailed descriptions of the datasets can be found in *SI Appendix*, Table S1. Moreover, we acquire 2,162 human housekeeping genes from the HRT Atlas (38) (https://housekeeping.unicamp.br) and 15 additional housekeeping genes from previous studies (39, 40). SpaCEX is publicly available at at https://github.com/WLatSunLab/SpaCEX.

## ACKNOWLEDGMENTS

The project is funded by Strategic Priority Research Program of Chinese Academy of Sciences (Grant No. XDB38050100) to H.W. X.S. is supported by Excellent Young Scientist Fund of Wuhan City (Grant No. 21129040740). We also thank Jiadi Lv, Daoli Wang, Suoya Han, Siyu Chen and Yuwei Hu for their helps in plotting figures and participation in discussions.

## Notes

https://singlecell.broadinstitute.org/single_cell/study/SCP815/sensitive-spatial-genome-wide-expression-profiling-at-cellular-resolution#study-summary

https://support.10xgenomics.com/spatial-gene-expression/datasets/1.1.0/V1_Breast_Cancer_Block_A_Section_1

https://www.ncbi.nlm.nih.gov/geo/query/acc.cgi?acc=GSM6801751

https://www.ncbi.nlm.nih.gov/geo/query/acc.cgi?acc=GSE178636

https://crukci.shinyapps.io/SpatialMouseAtlas/

https://housekeeping.unicamp.br

### Competing Interest Statement

The authors have declared no competing interest.

