## Supporting text, Figures S1 to S5, Tables S1 to S6, SI References for "Learning context-aware, distributed gene representations in spatial transcriptomics with SpaCEX"


**This PDF file includes:**

Supporting text
Figures S1 to S5
Tables S1 to S6
SI References

### Supporting Information Text

**Calculating group closeness centrality of gene clusters.** The group closeness centrality is a metric originally developed for quantifying the "closeness" of a group of nodes to other nodes on a graph(1). It can be used to measure the aggregated expression similarity between a gene cluster and other gene clusters on a weighted gene graph, with nodes representing genes and edge weights representing the expression dissimilarity between a pair of genes. In this study, we first convert the gene-to-gene similarity matrix $\mathbf{S} \in \mathrm{R}^{d\times d}$ of equation 2 (see "*Co-function analysis of genes within the same cluster*") into a distance matrix $\mathbf{D} \in \mathrm{R}^{d\times d}$:

$$\mathbf{D} = \mathrm{M} \times \mathrm{Diag}(\mathbf{S}) + \mathrm{Diag}(\mathbf{S}) \times \mathrm{M} - 2\mathbf{S}$$

Here, $\mathrm{M} \in \mathrm{R}^{d\times d}$ is a matrix full of 1s. Based on $\mathbf{D}$, the group closeness centrality of a gene cluster G, gcc(G), is calculated as:

$$\mathrm{gcc(G)} = \frac{|V - \mathrm{G}|}{\sum_{v\in\{V-\mathrm{G}\}} d_{\mathrm{G},v}}$$

$$\mathrm{d_{G,v}} = \mathrm{min}_{\mathrm{u}\in\{\mathrm{G}\}}(\mathbf{D}_{\mathrm{u,v}})$$

Here, $V$ represents the entire set of gene nodes, $|V - G|$ represents the number of gene nodes not belonging to $G$.

**Visualization of the aggregated gene expression pattern of a gene cluster.** To depict the collective gene expression pattern of a target gene cluster on the spatial map, we employ the "AddModuleScore" function from the Seurat R package to calculate module scores for the cluster across all spatial spots(2). Specifically, we bin all genes based on their average expression. Then a number of control genes are randomly selected from each bin to match the number of genes of the target gene cluster that reside in the same bin. Consequently, selected control genes should have a similar expression distribution to the genes within the target cluster but be functionally unrelated. The differences between the average expressions of the set of cluster genes and the set of control genes serve as the cluster's module scores across all spatial spots. Finally, a color spectrum is utilized to display module scores on the spatial map, with brighter hues signifying higher module scores, and vice versa.

**Biological analysis of SpaCEX-identified co-expressed gene clusters.** To investigate the biological processes implied by C1-C4, we perform GO enrichment analysis on their member genes to identify the 20 most significantly enriched GOBPs for each cluster. Additionally, we find that C1 (17 GOBPs) and C2 (18 GOBPs) are enriched in cancer related GOBPs (Table S6). Intriguingly, C1 contains more invasive cancer related GOBPs than C2, consistent with the observation that C1's aggregated gene expression overlaps with tissue regions abundant in IDCs. On the other hand, C3 contains fewer cancer-related but more benign stroma related GOBPs. Indeed, C3 member genes such as COL6A2 and COL1A2 in C3 are more involved in creating the extracellular matrix of benign stroma. In contrast, C4 displays no significant enrichment in GOBPs. Furthermore, to evaluate gene cofunctional coherence within the three clusters, we examine whether the involvement of a gene in a significantly enriched GOBP can be reliably predicated based on other member genes' involvement in the same GOBP (see *"Cofunction analysis of intra-cluster genes"*). The predictive accuracy is quantified by the area under the ROC curve (AUC) metric. Our results reveal that the predictions in C1 through C3 exhibit greater precision than those in C4, as evidenced by their superior AUC scores on the y-axis in *Main Text* Fig. 2*C*. Therefore, SpaCEX is capable of identifying clusters of cofunctional genes that are biologically or pathologically relevant to the dataset under investigation.

**Predicting gene-gene interactions using gene embeddings.** In this analysis, we adopt the methodology proposed by Du et al(3). A pair of genes is considered interacting (positive) if they share GO terms obtained using the R package "org.Hs.eg.db"; otherwise, they are considered noninteracting (negative). To reduce the number of false positive gene pairs, we omit the highly over-represented GO terms including "single transduction" (GO:0007165), three terms related to phosphorylation ("protein amino acid phosphorylation", GO:0006468; "protein amino acid autophosphorylation", GO:0046777; "protein amino acid dephosphorylation", GO:0006470), as well as all terms at the first three layers of the GO hierarchy. The universal gene set of this experiment is defined by the intersection of genes present in the hDLPFC dataset as well as the GO database, culminating in a total of 14,618 genes. Within this set, there are 778,293 positive samples involving 8,600 human genes, and 37,577,368 negative samples involving 8,759 human genes. All positive samples are included in the positive dataset, and an equal number of negative samples are randomly selected to constitute the negative dataset. Subsequently, 2% gene pairs are sampled from both the positive and negative datasets to create the training dataset, while the remaining 98% constitute the testing dataset. A gene-gene interaction predictor neural network (GGIPNN)(3), which takes gene embeddings as inputs, is employed to predict gene-gene interactions. *Main Text*, Fig. 3*C* illustrates that the GGIPNN, when using SGEs, achieves higher accuracy ($0.779 \pm 0.0048$) and AUC (0.831) scores than those using scBERT gene embeddings (accuracy=$0.742 \pm 0.0052$, AUC=0.812), Gene2vec gene embeddings (accuracy=$0.726 \pm 0.0026$, AUC=0.794), and random embeddings (accuracy=$0.500 \pm 0.0014$, AUC=0.510). Additionally, we introduce a bicolor heatmap to visualize the interactions among the top 1000 most interactive genes, with filled cells denoting gene interactions. *Main Text*, Fig. 3*D* showcases that the heatmap of predictions with SGEs exhibits greater concordance with the ground truth than those with others.

**Deriving the seeding gene-gene similarity matrix.** During the training of the model, an initial gene-gene similarity matrix is incorporated into the loss function $\mathcal{L}_1$ & $\mathcal{L}_2$ as a regularization term to inform the initial training phase of the model. We leverage multiple image recognition operators to extract feature descriptors from images of gene spatial expression maps, based on which the seeding gene-gene similarity matrix is calculated. Specifically, on the gray-scale level of the image, we utilize Sober operator to extract gradient magnitude and orientation descriptors, Laplacian operator to extract gradient divergence descriptor, and Canny operators to extract the gradient continuity descriptor. Meanwhile, three average and standard deviation pooling filters of different sizes are used to extract patch brightness descriptors. The normalized spatial expression matrix of gene $u$ is denoted as $\mathrm{X}^u \in \mathbb{R}^{N_x \times N_y}$, where $N_x$ *and* $N_y$ denote the number of spatial spots along the horizontal and vertical directions of the spatial map. Out-of-tissue spatial spots are all padded with 0s. $\mathrm{X}^u$ is smoothened with a convolutional Gaussian kernel $\mathrm{H} \in \mathcal{R}^{d \times \mathrm{d}}$, obtaining a denoised expression matrix $\tilde{X}^u$:

$$H_{i,j} = \frac{1}{2\pi\sigma^2} exp\left(-\frac{(i-(k-1)/2)^2+(j-(k-1)/2)^2}{2\sigma^2}\right); 1 \le i,j \le d;$$

$$\tilde{X}^u_{i,j} = sum\left(H \odot X^u_{s(k),t(k)}\right);$$

Next, Sober, Laplacian and Canny operators are applied on $\tilde{X}^u$ to generate matrices of corresponding descriptors. For a specific gene $u$, let $\mathcal{G}^u$ and $\Theta^u$ denote the matrices of gradient magnitude and orientation descriptors, $\mathcal{L}^u$ the matrix of gradient divergence descriptor, and $\mathcal{C}^u$ the matrix of gradient continuity descriptor. The pooling filters are applied by segmenting $\tilde{X}^u$ into small patches containing $k \times k$ *spots, where* $k \in \{1,3,5\}$, from which three patch brightness mean matrices $\mathcal{A}^u(k)$ and two variance matrices $\mathcal{S}^u(k)$ (eligible only for $k = 3,5$) are calculated:

$$\mathcal{A}^u_{i,j}(k) = \mathrm{avg}\left(X^u_{s(k),t(k)}\right),$$

$$\mathcal{S}^u_{i,j}(k) = \mathrm{std}\left(X^u_{s(k),t(k)}\right),$$

$$1 \le i \le N_x, 1 \le j \le N_y$$

$$\mathrm{s(k)} = \left[i - \frac{(k-1)}{2} : i + \frac{(k-1)}{2}\right], \mathrm{t(k)} = \left[j - \frac{(k-1)}{2} : j + \frac{(k-1)}{2}\right]$$

Finally, the initial gene similarity matrix S is calculated as the average Pearson correlation between gene pairs' descriptor matrices:

$$\mathrm{S}_{u,v} = \begin{cases} \mathrm{avg}\left(\rho_{u,v}(\Xi^u, \Xi^v)\right); \Xi \in \{\mathcal{A}(k), \mathcal{S}(k), \mathcal{G}, \Theta, \mathcal{L}, \mathcal{C}\},\ \mathrm{k} \in 1,3,5, & if\ \mathrm{u} \neq \mathrm{v} \\ 0, \quad if\ u = v \end{cases}$$

Here, $u, v$ represent any two genes in the dataset.

**MAP-EM inference of SMM parameters.** We first list the mathematical notations used in the inference below:

---

$X \in \mathcal{R}^{N \times M}$: the raw gene count matrix.
$N$: the number of genes.
$M$: the number of spatial spots.
$K$: the number of gene clusters (components in the SMM).
$Z \in \mathcal{R}^{N \times D} = f_{encoder}(W, X)$: the SpaCEX-generated gene embeddings.
$W$: parameters of the MAE encoder.
$\hat{X} \in \mathcal{R}^{N \times M} = f_{decoder}(\widehat{W}, Z)$: the reconstructed gene count matrix by the decoder.

$\widehat{W}$: parameters of the MAE decoder.
$\phi_k(\mu_k, \Sigma_k, v_k)$: the $pdf$ of the $k$-th SMM component.
$\Pi = \{\pi_k, \forall k \in [1, K]\}$: the weights of SMM components.
$\Theta = \{\theta_k: \mu_k, \Sigma_k, v_k, \pi_k, \forall k \in [1, K]\}$: parameters of the SMM components.
$\xi_i \in [1, K]$: the SMM component membership of $x_i$.

---

We re-write the Student's t distribution of the $k$-th component of the SMM as a Gaussian scale mixture:

$$\phi(z_i|\mu_k, \Sigma_k, v_k) = \int N\left(z_i\middle|\mu_k, \frac{\Sigma_k}{\zeta_{i,k}}\right)\Gamma\left(\zeta_{i,k}\middle|\frac{v_k}{2}, \frac{v_k}{2}\right)d\zeta_{i,k}$$

Therefore, under the EM framework, all hidden variables are $H = \{h_i: \xi_i, \zeta_{i,k}, \forall i \in [1, N], \forall k \in [1, K]\}$. The complete data log likelihood is:

$$\ell_c(\Theta) = \log P(X, H|\Theta) = \sum_i \sum_k II(\xi_i = k)\left(log\pi_k + logf\left(z_i, \zeta_{i,k}\middle|\mu_k, \Sigma_k, v_k\right)\right)$$

$$logf\left(z_i, \zeta_{i,k}\middle|\mu_k, \Sigma_k, v_k\right) \propto log\Gamma\left(\zeta_{i,k}\middle|\frac{v_k}{2}, \frac{v_k}{2}\right) + \frac{D}{2}log\zeta_{i,k} - \frac{1}{2}log|\Sigma_k| - \frac{1}{2}\zeta_{i,k}\sigma_{i,k}$$

$$\sigma_{i,k} = (z_i - \mu_k)^T\Sigma_k^{-1}(z_i - \mu_k)$$

E step. In the $t$-th iteration, we have the auxiliary function $Q$ as:

$$Q\left(\Theta, \Theta^{(t-1)}\right) = E\left(\ell_c(\Theta)\middle|\Theta^{(t-1)}\right)$$

$$= \sum_i \sum_k p\left(\xi_i = k\middle|z_i, \Theta^{(t-1)}\right)\left(log\pi_k^{(t-1)} + E\left(logf\left(z_i, \zeta_{i,k}\middle|\mu_k^{(t-1)}, \Sigma_k^{(t-1)}, v_k^{(t-1)}\right)\right)\right).$$

The expected sufficient statistics (ESS) are:

$$\overline{\xi_{i,k}}^{(t)} = p\left(\xi_i = k\middle|z_i, \Theta^{(t-1)}\right) = \frac{\pi_k^{(t-1)}\phi\left(z_i|\mu_k^{(t-1)}, \Sigma_k^{(t-1)}, v_k^{(t-1)}\right)}{\sum_{k'}\pi_{k'}^{(t-1)}\phi\left(z_i|\mu_{k'}^{(t-1)}, \Sigma_{k'}^{(t-1)}, v_{k'}^{(t-1)}\right)}$$

$$\overline{\zeta_{i,k}}^{(t)} = E\left(p\left(\zeta_{i,k}\middle|z_i, \theta_k^{(t-1)}\right)\right) = E\left(\Gamma\left(\zeta_{i,k}\middle|\frac{v_k^{(t-1)} + D}{2}, \frac{v_k^{(t-1)} + \sigma_{i,k}^{(t-1)}}{2}\right)\right) = \frac{v_k^{(t-1)} + D}{v_k^{(t-1)} + \sigma_{i,k}^{(t-1)}}$$

Then the complete data log likelihood of $z_i$ becomes:

$$E\left(logf\left(z_i, \zeta_{i,k}\middle|\mu_k, \Sigma_k, v_k\right)\right) \propto G(z_i, \mu_k, \Sigma_k)^{(t)} + F\left(\zeta_{i,k}, v_k\right)^{(t)}, \quad (2)$$

$$G(z_i, \mu_k, \Sigma_k)^{(t)} = -\frac{1}{2}log|\Sigma_k| - \frac{\overline{\zeta_{i,k}}^{(t)}}{2}\sigma_{i,k}$$

$$F\left(\zeta_{i,k}, v_k\right)^{(t)} = \frac{v_k log(v_k/2)}{2} - \Gamma\left(\frac{v_k}{2}\right) + \frac{v_k}{2}\left(\overline{log\zeta_{i,k}}^{(t)} - \overline{\zeta_{i,k}}^{(t)}\right).$$

M step. In the $t$-th iteration, we maximize $Q$ with respect to $\forall\theta_k \in \Theta$. Rather than achieving the MLE, we introduce a prior distribution on $\theta_k$ and solve for MAP of $\theta_k$ to alleviate model overfitting. Specifically, we introduce a conjugate prior on $\Pi$ as a Dirichlet distribution, and a conjugate prior on $\{\mu_k, \Sigma_k\}$ as a normal-inverse Wishart (NIW) distribution:

$$\mathrm{Dir}(\Pi|\alpha^0) \equiv \frac{1}{\mathrm{B}(\alpha^0)}\prod_k \pi_k^{\alpha_k^0 - 1},$$

$$\mathrm{NIW}(\mu_k, \Sigma_k|m_0, \kappa_0, S_0, \rho_0)$$

$$\propto |\Sigma_k|^{-\frac{1}{2}}exp\left(-\frac{\kappa_0}{2}(\mu_k - m_0)^T\Sigma_k^{(-1)}(\mu_k - m_0)\right) \times |\Sigma_k|^{-\frac{(\rho_0 + D + 1)}{2}}\exp\left(-\frac{1}{2}tr(S_0\Sigma_k^{-1})\right).$$

Here, we have weaker priors as $\alpha^0 = \vec{1}$, $\rho_0 = D + 2$, $S_0 = N^{-1}diag\left(\left(Z - \frac{1}{N}\Omega Z\right)^T\left(Z - \frac{1}{N}\Omega Z\right)\right)$, $\Omega = \vec{1}^T \times \vec{1}$, $\kappa_0 = 0$, $m_0 = \frac{\sum_i z_i}{N}$. The posterior distribution of $\Pi$ and $\{\mu_k, \Sigma_k\}$ are:

$$p(\Pi|X) \sim \mathrm{Dir}\left(\Pi\middle|\alpha^{(t)}\right) \equiv \frac{1}{\mathrm{B}(\alpha^{(t)})}\prod_k \pi_k^{\alpha_k^{(t)} - 1}$$

$$\alpha_k^{(t)} = \alpha_k^0 + \sum_i \overline{\xi_{i,k}}^{(t)}, \forall k \in [1, K]$$

$$\mathrm{p}(\mu_k,\Sigma_k|X)\sim NIW\left(\mu_k,\Sigma_k\middle|\mathrm{m}_k^{(t)},\kappa_k^{(t)},S_k^{(t)},\rho_k^{(t)}\right)$$
$$\kappa_k^{(t)}=\kappa_0+\overline{\omega_k}^{(t)}=\overline{\omega_k}^{(t)}$$
$$\overline{\omega_{i,k}}^{(t)}=\overline{\xi_{i,k}}^{(t)}\overline{\zeta_{i,k}}^{(t)}$$
$$\overline{\omega_k}^{(t)}=\sum_i\overline{\omega_{i,k}}^{(t)}$$
$$\rho_k^{(t)}=\rho_0+\overline{\xi_k}^{(t)}$$
$$\overline{\xi_k}^{(t)}=\sum_i\overline{\xi_{i,k}}^{(t)}$$
$$\mathrm{m}_k^{(t)}=\frac{\overline{\omega_k}^{(t)}\overline{z}_k^{(t)}+\kappa_0 m_0}{\kappa_k^{(t)}}=\overline{z}_k^{(t)}$$
$$\overline{z}_k^{(t)}=\frac{\sum_i\left(\overline{\omega_{i,k}}^{(t)}z_i\right)}{\overline{\omega_k}^{(t)}}$$
$$S_k^{(t)}=S_0+\sum_i\overline{\omega_{i,k}}^{(t)}z_iz_i^T+\kappa_0m_0m_0^T-\kappa_k^{(t)}\mathrm{m}_k^{(t)}\left(\mathrm{m}_k^{(t)}\right)^T$$
$$=S_0+\sum_i\overline{\omega_{i,k}}^{(t)}z_iz_i^T-\kappa_k^{(t)}\mathrm{m}_k^{(t)}\left(\mathrm{m}_k^{(t)}\right)^T.$$

Then we have the MAP estimates of $\pi_k$ and $\{\mu_k,\Sigma_k\}$ as $\pi_k^{(t)}$ and $\{\mu_k^{(t)},\Sigma_k^{(t)}\}$:

$$\pi_k^{(t)}=\frac{\alpha_{\mathrm{k}}^{(t)}-1}{\sum_{k'}\alpha_{k'}^{(t)}-K}$$
$$\mu_k^{(t)}=\mathrm{m}_k^{(t)}=\overline{z}_k^{(t)}$$
$$S_k^{(t)}=S_0+\sum_i\left[\overline{\omega_{i,k}}^{(t)}\left(z_i-\mathrm{m}_k^{(t)}\right)\left(z_i-\mathrm{m}_k^{(t)}\right)^T\right]=S_0+S_{\mathrm{mle},k}^{(t)}$$
$$S_{\mathrm{mle},k}^{(t)}=\sum_i\left[\overline{\omega_{i,k}}^{(t)}\left(z_i-\overline{z}_k^{(t)}\right)\left(z_i-\overline{z}_k^{(t)}\right)^T\right]$$
$$\Sigma_k^{(t)}=\frac{S_k^{(t)}}{\rho_k^{(t)}+D+2}=\frac{S_0+S_{\mathrm{mle},k}^{(t)}}{\widehat{\rho_0}+\overline{\xi_k}^{(t)}}=\frac{\widehat{\rho_0}}{\widehat{\rho_0}+\overline{\xi_k}^{(t)}}\cdot\frac{S_0}{\widehat{\rho_0}}+\frac{\overline{\xi_k}^{(t)}}{\widehat{\rho_0}+\overline{\xi_k}^{(t)}}\cdot\frac{S_{\mathrm{mle},k}^{(t)}}{\overline{\xi_k}^{(t)}}=\beta\Sigma_0+(1-\beta)\Sigma_{mle,\mathrm{k}}^{(t)}.$$

Then we have:

$$\Sigma_k^{(t)}(i,j)=\begin{cases}\beta\Sigma_0(i,j)+(1-\beta)\Sigma_{mle,\mathrm{k}}^{(t)}(i,j), if\ i=j\\(1-\beta)\Sigma_{mle,\mathrm{k}}^{(t)}(i,j), otherwise\end{cases}.$$

The off-diagonal entries in $\Sigma_k^{(t)}$ are shrunk toward 0 to promote its sparsity, thereby reducing the computational load and possibility of overfitting.

$v_k^{(t)}$ can be derived by maximizing $\sum_{\mathrm{i}}\left(\overline{\xi_{i,k}}^{(t)}\cdot F\left(\zeta_{i,k}^{(t)},v_k^{(t)}\right)\right)$. However, there is no closed-form solution, so we apply the generalized EM (GEM) to approximate the solution as follows:

$$\zeta_{i,k}^{(t)}\sim\Gamma\left(\zeta_{i,k}^{(t)}\middle|\frac{v_k^{(t-1)}+D}{2},\frac{v_k^{(t-1)}+\sigma_{i,k}^{(t-1)}}{2}\right)$$
$$\Rightarrow\overline{log\zeta_{i,k}}^{(t)}=E\left(log\zeta_{i,k}^{(t)}\right)=\Psi\left(\frac{v_k^{(t-1)}+D}{2}\right)-log\frac{v_k^{(t-1)}+\sigma_{i,k}^{(t-1)}}{2},$$

where $\Psi(\mathrm{x})\equiv\frac{\mathrm{d}}{\mathrm{dx}}\log\Gamma(\mathrm{x})$ is the digamma function. Then we have:

$$\frac{\mathrm{d}}{\mathrm{d}v_k^{(t)}}\sum_{\mathrm{i}}\left(\overline{\xi_{i,k}}^{(t)}\cdot F\left(\zeta_{i,k}^{(t)},v_k^{(t)}\right)\right)=\sum_{\mathrm{i}}\overline{\xi_{i,k}}^{(t)}\left(\frac{1}{2}\log\left(\frac{v_k^{(t)}}{2}\right)+\frac{1}{2}-\frac{1}{2}\Psi\left(\frac{v_k^{(t)}}{2}\right)+\frac{1}{2}\left(\overline{log\zeta_{i,k}}^{(t)}-\overline{\zeta_{i,k}}^{(t)}\right)\right),$$

Then $v_k^{(t)} \leftarrow v_k^{(t)} - \lambda \cdot \frac{\mathrm{d}}{\mathrm{d}v_k^{(t)}} \mathrm{F}\left(\zeta_{i,k}^{(t)}, v_k^{(t)}\right)$ is repeated for several times to achieve a "partial" improvement to $v_k^{(t)}$, which still guarantees the convergence to a local optimum. Next, the EM algorithm continues to E step of the $(t+1)-th$ iteration to update $H^{(t+1)} = \{h_{i,k}^{(t+1)}: \overline{\xi_{i,k}}^{(t+1)}, \overline{\zeta_{i,k}}^{(t+1)}, \forall i \in [1,N], \forall k \in [1,K]\}$ until either convergence is achieved, or a pre-specified number of iterations is reached. Finally, the score and soft assignment of $z_i$ to the $k$-th component ($\mathcal{q}_{i,k}$) can be calculated by plugin $\theta_k$ as:

$$q_{i,k} = \pi_k \phi(z_i | \mu_k, \Sigma_k, v_k)$$
$$\mathcal{q}_{i,k} = \frac{q_{i,k}}{\sum_j q_{i,j}}, \forall i \in [1,N], \forall k \in [1,K]$$

**The joint optimization of the gene representation learning model and SMM via a discriminative boosted clustering.** In this section, we focus on deriving the gradients of $\mathcal{L}_{\ell\ell}$ $and$ $\mathcal{L}_{size}$ with respect to $Z$, and the gradients of $\mathcal{L}_c$ with respect to $Z$ and $\Theta$. The derivations of the gradients of $\mathcal{Lap\ell}$ and $\mathcal{L}_r$ with respect to $Z$ and the gradient of $\mathcal{L}_r$ with respect to $\hat{X}$ are relatively trivial and therefore ignored.

$$\mathcal{L}_{\ell\ell} = \log\mathcal{P}(Z|\Theta) = \sum_{\mathrm{i}=1}^{\mathrm{N}} \log\left[\sum_k q_{i,k}\right]$$
$$\mathcal{L}_c = KL(\mathcal{P}|\mathcal{Q}) = \sum_i^N \sum_j^K \mathcal{p}_{i,j} \, log \frac{\mathcal{p}_{i,j}}{\mathcal{q}_{i,j}}$$
$$\mathcal{p}_{i,k} = \frac{\mathcal{q}_{i,k}^2 / \sum_i \mathcal{q}_{i,j}}{\sum_j \left(\mathcal{q}_{i,j}^2 / \sum_i \mathcal{q}_{i,j}\right)}$$
$$\mathcal{L}_{size}(Z,\Theta) = \sum_{k=1}^{K} -J_k \log J_k$$

where $J_k = \begin{cases} \frac{\sum_i^N \mathcal{q}_{i,k}}{N}, & if\ J_k \leq \tau \\ 1, otherwise \end{cases}$

The density function of $z_i$ given $\{\mu_k, \Sigma_k, v_k\}$ is:

$$\phi(z_i|\mu_k,\Sigma_k,v_k) \propto \frac{\Gamma\left(\frac{v_k+\mathrm{D}}{2}\right)}{\Gamma\left(\frac{v_k}{2}\right)} v_k^{-\frac{D}{2}} |\Sigma_k|^{-\frac{1}{2}} \left[1 + \frac{1}{v_k}(z_i - \mu_k)^{\mathrm{T}} \Sigma_k^{-1} (z_i - \mu_k)\right]^{-\left(\frac{v_k+\mathrm{D}}{2}\right)}$$
$$= \hbar(v_k)|\Sigma_k|^{-\frac{1}{2}} \left[1 + \frac{\sigma_{i,k}}{v_k}\right]^{-\left(\frac{v_k+\mathrm{D}}{2}\right)} = \hbar(v_k)|\Sigma_k|^{-\frac{1}{2}} \mathrm{u}_{\mathrm{i},k}^{-\left(\frac{v_k+\mathrm{D}}{2}\right)}$$

$$\frac{\partial \mathrm{u}_{\mathrm{i},k}}{\partial z_i} = \frac{2}{v_k} \Sigma_k^{-1}(z_i - \mu_k) \Longrightarrow$$
$$\frac{\partial q_{i,k}}{\partial z_i} = \frac{\partial q_{i,k}}{\partial \mathrm{u}_{\mathrm{i,k}}} \cdot \frac{\partial \mathrm{u}_{\mathrm{i},k}}{\partial z_i} = -\pi_k \hbar(v_k)|\Sigma_k|^{-\frac{1}{2}} \left(\frac{v_k+\mathrm{D}}{2}\right) \mathrm{u}_{\mathrm{i},k}^{-\left(\frac{v_k+\mathrm{D}}{2}+1\right)} \cdot \frac{2}{v_k} \Sigma_k^{-1}(z_i - \mu_k)$$
$$= -\left(\frac{v_k+\mathrm{D}}{v_k}\right) \mathrm{u}_{\mathrm{i,k}}^{-1} q_{i,k} \cdot \Sigma_k^{-1}(z_i - \mu_k)$$

$$\frac{\partial q_{i,j}}{\partial \mu_{\mathrm{k}}} = \begin{cases} \frac{\partial q_{i,k}}{\partial \mathrm{u}_{\mathrm{i},k}} \cdot \frac{\partial \mathrm{u}_{\mathrm{i},k}}{\partial \mu_k} = -\left(\frac{v_k+\mathrm{D}}{v_k}\right) \mathrm{u}_{\mathrm{i,k}}^{-1} q_{i,k} \cdot \Sigma_k^{-1}(\mu_k - z_i), \mathrm{j} = k \\ 0, \mathrm{j} \neq k \end{cases}$$

$$\frac{\partial q_{i,j}}{\partial \Sigma_k} = \begin{cases} \frac{\partial q_{i,k}}{\partial \mathrm{u_{i,k}}} \cdot \frac{\partial \mathrm{u_{i,k}}}{\partial \Sigma_k} + \frac{\partial q_{i,k}}{\partial |\Sigma_k|} \cdot \frac{\partial |\Sigma_k|}{\partial \Sigma_k}, \mathrm{j} = k \\ 0, \mathrm{j} \neq k \end{cases}$$

$$= \begin{cases} q_{i,k}\left(\left(\frac{v_k + \mathrm{D}}{2v_k}\right) \mathrm{u_{i,k}^{-1}} \cdot \ \Sigma_k^{-1}(z_i - \mathrm{\mu_k})(z_i - \mathrm{\mu_k})^{\mathrm{T}} \Sigma_k^{-1} - \frac{1}{2}\ \Sigma_k^{-1}\right) = q_{i,k}\mathscr{f}(z_i, \mathrm{\mu}_k, v_k, \Sigma_k, D), \mathrm{j} = k \\ 0, \mathrm{j} \neq k \end{cases}$$

$$\frac{\partial q_{i,j}}{\partial v_k} = \begin{cases} q_{i,k}\frac{\partial ln(q_{i,k})}{\partial v_k}, \mathrm{j} = k \\ 0, \mathrm{j} \neq k \end{cases}$$

$$= \begin{cases} q_{i,k}\left(\frac{v_k + D}{2}\mathrm{u_{i,k}^{-1}}\frac{\sigma_{i,k}}{v_k^2} - \frac{1}{2}ln u_{i,k} + \frac{1}{2}\Gamma\left(\frac{v_k + D}{2}\right)\Psi\left(\frac{v_k + \mathrm{D}}{2}\right) - \frac{1}{2}\Gamma\left(\frac{v_k}{2}\right)\Psi\left(\frac{v_k}{2}\right) - \frac{\mathrm{D}}{2v_k}\right), \mathrm{j} = k \\ 0, \mathrm{j} \neq k \end{cases}$$

$$= \begin{cases} q_{i,k}\mathscr{g}(z_i, \mathrm{\mu}_k, v_k, \Sigma_k, D), \mathrm{j} = k \\ 0, \mathrm{j} \neq k \end{cases}$$

$$\frac{\partial q_{i,j}}{\partial \pi_k} = \begin{cases} \frac{q_{i,k}}{\pi_k}, \mathrm{j} = k \\ 0, \mathrm{j} \neq k \end{cases}$$

By chain rules of derivatives, we have:

$$\frac{\partial \mathscr{q}_{i,k}}{\partial z_i} = \frac{\frac{\partial q_{i,k}}{\partial z_i} \cdot \sum_j q_{i,j} - q_{i,k} \cdot \sum_j \frac{\partial q_{i,j}}{\partial z_i}}{\left(\sum_j q_{i,j}\right)^2}$$

$$= -\left(\frac{\mathrm{v}_k + \mathrm{D}}{\mathrm{v}_k}\right)\mathscr{q}_{i,k}\left(\mathrm{u_{i,k}^{-1}} \cdot \ \Sigma_k^{-1}(z_i - \mathrm{\mu_k}) - \sum_{\mathrm{j}} \mathrm{u_{i,j}^{-1}}\,\mathscr{q}_{i,j} \cdot \ \Sigma_j^{-1}(z_i - \mathrm{\mu_j})\right)$$

$$\frac{\partial \mathscr{q}_{i,j}}{\partial \mathrm{\mu_k}} = \begin{cases} \frac{\frac{\partial q_{i,j}}{\partial \mathrm{\mu_k}} \cdot \sum_j q_{i,j} - q_{i,j} \cdot \sum_j \frac{\partial q_{i,j}}{\partial \mathrm{\mu_k}}}{\left(\sum_j q_{i,j}\right)^2}, \ \ \mathrm{j} \neq k \\ \frac{\frac{\partial q_{i,k}}{\partial \mathrm{\mu_k}} \cdot \sum_j q_{i,j} - q_{i,k} \cdot \sum_j \frac{\partial q_{i,j}}{\partial \mathrm{\mu_k}}}{\left(\sum_j q_{i,j}\right)^2}, \mathrm{j} = k \end{cases}$$

$$= \begin{cases} \left(\frac{\mathrm{v}_k + \mathrm{D}}{\mathrm{v}_k}\right)\mathscr{q}_{i,j}\left(\mathrm{u_{i,k}^{-1}}\mathscr{q}_{i,k} \cdot \ \Sigma_k^{-1}(\mathrm{\mu_k} - z_i)\right), \ \ \mathrm{j} \neq k \\ -\left(\frac{\mathrm{v}_k + \mathrm{D}}{\mathrm{v}_k}\right)\mathscr{q}_{i,k}\left(\mathrm{u_{i,k}^{-1}} \cdot \ \Sigma_k^{-1}(\mathrm{\mu_k} - z_i) - \mathrm{u_{i,k}^{-1}}\mathscr{q}_{i,k} \cdot \ \Sigma_k^{-1}(\mathrm{\mu_k} - z_i)\right), \mathrm{j} = k \end{cases}$$

$$\frac{\partial \mathscr{q}_{i,j}}{\partial \Sigma_k} = \begin{cases} \frac{\frac{\partial q_{i,j}}{\partial \Sigma_k} \cdot \sum_j q_{i,j} - q_{i,j} \cdot \sum_j \frac{\partial q_{i,j}}{\partial \Sigma_k}}{\left(\sum_j q_{i,j}\right)^2}, \ \ \mathrm{j} \neq k \\ \frac{\frac{\partial q_{i,k}}{\partial \Sigma_k} \cdot \sum_j q_{i,j} - q_{i,k} \cdot \sum_j \frac{\partial q_{i,j}}{\partial \Sigma_k}}{\left(\sum_j q_{i,j}\right)^2}, \ \ \mathrm{j} = k \end{cases}$$

$$= \begin{cases} -\mathscr{q}_{i,j}\left(\mathscr{q}_{i,k} \cdot \mathscr{f}(z_i, \mathrm{\mu}_k, v_k, \Sigma_k, D)\right), \mathrm{j} \neq k \\ \mathscr{q}_{i,k}\left(\mathscr{f}(z_i, \mathrm{\mu}_k, v_k, \Sigma_k, D) - \mathscr{q}_{i,k} \cdot \mathscr{f}(z_i, \mathrm{\mu}_k, v_k, \Sigma_k, D)\right), \mathrm{j} = k \end{cases}$$

$$\frac{\partial q_{i,j}}{\partial v_k}=\begin{cases}\dfrac{\frac{\partial q_{i,j}}{\partial v_k}\cdot\sum_j q_{i,j}-q_{i,j}\cdot\sum_j\frac{\partial q_{i,j}}{\partial v_k}}{\left(\sum_j q_{i,j}\right)^2}=-q_{i,j}q_{i,k}g(z_i,\mu_k,v_k,\Sigma_k,D),\ \mathrm{j}\neq k\\ \dfrac{\frac{\partial q_{i,k}}{\partial v_k}\cdot\sum_j q_{i,j}-q_{i,k}\cdot\sum_j\frac{\partial q_{i,j}}{\partial v_k}}{\left(\sum_j q_{i,j}\right)^2}=q_{i,k}\left(1-q_{i,k}\right)g(z_i,\mu_k,v_k,\Sigma_k,D),\ \mathrm{j}=k\end{cases}$$

$$\frac{\partial q_{i,j}}{\partial \pi_k}=\begin{cases}\dfrac{\frac{\partial q_{i,j}}{\partial \pi_k}\cdot\sum_j q_{i,j}-q_{i,j}\cdot\sum_j\frac{\partial q_{i,j}}{\partial \pi_k}}{\left(\sum_j q_{i,j}\right)^2}=-\dfrac{q_{i,j}q_{i,k}}{\pi_k},\mathrm{j}\neq k\\ \dfrac{\frac{\partial q_{i,k}}{\partial \pi_k}\cdot\sum_j q_{i,j}-q_{i,k}\cdot\sum_j\frac{\partial q_{i,j}}{\partial \pi_k}}{\left(\sum_j q_{i,j}\right)^2}=\dfrac{q_{i,k}\left(1-q_{i,k}\right)}{\pi_k},\mathrm{j}=k\end{cases}$$

Then the derivatives of $\mathcal{L}_{\ell\ell}$ and $\mathcal{L}_{size}$ with respect to $z_i$ are:

$$\frac{\partial\mathcal{L}_{\ell\ell}}{\partial z_i}=\frac{\sum_j\frac{\partial q_{i,j}}{\partial z_i}}{\sum_j q_{i,j}}=-\left(\frac{v_k+\mathrm{D}}{v_k}\right)\frac{\mathrm{u}_{\mathrm{i,k}}^{-1}q_{i,k}\cdot\Sigma_k^{-1}(z_i-\mu_k)}{\sum_j q_{i,j}}$$

$$\frac{\partial\mathcal{L}_{size}}{\partial z_i}=\sum_{\mathrm{j}\in\{J_j\leq\tau\}}\frac{-(1+\log J_{\mathrm{j}})}{\mathrm{N}}\sum_{\mathrm{i}=1}^{\mathrm{N}}\frac{\partial q_{i,\mathrm{j}}}{\partial z_i}$$

As the target distribution $\mathcal{P}$ is fixed during the joint optimization within an epoch, we have the derivatives of $\mathcal{L}_c$ with respect to $z_i$ and $\theta_k=\{\mu_k,\Sigma_k,v_k,\pi_k\}$ as:

$$\frac{\partial\mathcal{L}_c}{\partial z_i}=-\sum_j\left[\frac{p_{i,j}}{q_{i,j}}\cdot\frac{\partial q_{i,j}}{\partial x_i}\right]=\left(\frac{v_k+\mathrm{D}}{v_k}\right)\sum_j\left(p_{i,k}-q_{i,k}\right)\mathrm{u}_{\mathrm{i,k}}^{-1}\cdot\Sigma_k^{-1}(z_i-\mu_k)$$

$$\frac{\partial\mathcal{L}_c}{\partial \mu_k}=-\sum_i\sum_j\left[\frac{p_{i,j}}{q_{i,j}}\cdot\frac{\partial q_{i,j}}{\partial \mu_k}\right]=\left(\frac{v_k+\mathrm{D}}{v_k}\right)\sum_i\left(p_{i,k}-q_{i,k}\right)\mathrm{u}_{\mathrm{i,k}}^{-1}\cdot\Sigma_k^{-1}(\mu_k-z_i)$$

$$\frac{\partial\mathcal{L}_c}{\partial \Sigma_k}=-\sum_i\sum_j\left[\frac{p_{i,j}}{q_{i,j}}\cdot\frac{\partial q_{i,j}}{\partial \Sigma_k}\right]=\sum_i\left(q_{i,k}-p_{i,k}\right)f(z_i,\mu_k,v_k,\Sigma_k,D)$$

$$\frac{\partial\mathcal{L}_c}{\partial v_k}=-\sum_i\sum_j\left[\frac{p_{i,j}}{q_{i,j}}\cdot\frac{\partial q_{i,j}}{\partial v_k}\right]=-\sum_i\left(p_{i,k}-q_{i,k}\right)g(z_i,\mu_k,v_k,\Sigma_k,D)$$

$$\frac{\partial\mathcal{L}_c}{\partial \pi_k}=-\sum_i\sum_j\left[\frac{p_{i,j}}{q_{i,j}}\cdot\frac{\partial q_{i,j}}{\partial \pi_k}\right]=-\sum_i\frac{\left(p_{i,k}-q_{i,k}\right)}{\pi_k}$$

**SpaCEX-SPS.** In this section, we introduce a generalized linear model (GLM)-based method, SpaCEX-spatial pattern simulator (SpaCEX-SPS), to simulate genes with desired spatial expression patterns, based on which real genes with similar expression patterns can be identified. This task is not trivial—on one hand, given no prior knowledge about which real genes exhibit the target expression pattern, existing reference-based simulation methods, as seen in Andersson et al(4)., fail to pinpoint an appropriate gene for reference. On the other hand, without referring to real target data, reference-free simulation methods like SRTsim(5) fall short of generating ST data with consistent data properties.

SpaCEX-SPS allows specifying precise gene expression levels in targeted tissue regions. Specifically, let $G$ represent the set of all genes and $S$ the set of all spots in the target dataset. The size-factor-normalized read counts of any gene $\mathrm{i}\in\mathrm{G}$ at spot $\mathrm{j}\in\mathrm{S}$, $\mathrm{X}_{\mathrm{i,j}}$, follows a negative binomial distribution:

$$X_{i,j} \sim NB(\mu_i, s_i),$$

where $\mu_i$ and $s_i$ are mean and dispersion parameters, respectively. Then we have the following equations:

$$\sigma_i^2 = \mu_i + \frac{\mu_i^2}{s_i} \Rightarrow cv_i^2 = \frac{1}{\mu_i} + \frac{1}{s_i}, \tag{2}$$

where $\sigma_i^2$ and $cv_i^2$ represent variance and squared coefficient of variation (SCV) of gene $i$, respectively. As proved in Brennecke et al(6)., we have:

$$E\left[\widehat{cv}_i^2\right] \approx a_0 + \frac{a_1}{\hat{\mu}_i},$$

$$\hat{\mu}_i = \frac{\sum_j X_{i,j}}{N},$$

$$\widehat{cv}_i^2 = \frac{\sum_j \left(X_{i,j} - \hat{\mu}_i\right)^2 / (N-1)}{\hat{\mu}_i^2},$$

where $\widehat{cv}_i^2$ and $\hat{\mu}_i$ are sample SCV and mean, respectively. $N$ denotes the total number of spots in the ST dataset. Note that $\widehat{cv}_i^2$ approximately follows a $\chi^2$ or gamma distribution. By taking $\frac{1}{\hat{\mu}_i}$ as input, we can use a GLM of the gamma family with an identity link function to perform the regression:

$$\sum_i \log\left(P\left(\widehat{cv}_i^2 \middle| \alpha, \beta_i\right)\right) = \sum_i \left(-\alpha\log(\beta_i) - \frac{\widehat{cv}_i^2}{\beta_i} + (\alpha - 1)\log\left(\widehat{cv}_i^2\right)\right) + C_1,$$

$$E\left[\widehat{cv}_i^2 \middle| \alpha, \beta_i\right] = \alpha\beta_i = \eta_i = a_0 + \frac{a_1}{\hat{\mu}_i},$$

where $\alpha = \frac{1}{\sigma}$ is a constant, and $\sigma$ denotes the GLM dispersion parameter. Let the natural parameter of the GLM be $\theta_i = \frac{-1}{E\left[\widehat{cv}_i^2 \middle| \alpha, \beta_i\right]}$, then we have:

$$\sum_i \log\left(P\left(\widehat{cv}_i^2 \middle| \alpha, \beta_i\right)\right) \propto \sum_i \left(\frac{\theta_i\widehat{cv}_i^2 - \log\left(-\frac{1}{\theta_i}\right)}{\sigma} + \frac{1-\sigma}{\sigma}\log\left(\widehat{cv}_i^2\right)\right),$$

from which coefficients $a_0$ and $a_1$ can be estimated using the Fisher-scoring method.

Assume we aim to simulate a gene with a top $t\%$ expression level in a spatial region $S_1 \in S$, and the gene conforms to a NB distribution with mean $\mu$ and dispersion $s$. We first calculate and rank ascendingly the average expressions of all genes in the target dataset, denoted as $L$. Then, $\mu$ and the corresponding squared coefficient of variation, cv, can be estimated as:

$$\mu = L@t\%,$$

$$cv^2 \approx E[cv^2] = \alpha_0 + \frac{\alpha_1}{\mu}.$$

According to equation 2, $s$ can be computed as:

$$s = \frac{1}{cv^2 - \frac{1}{\mu}}.$$

Let $N_1$ denote the number of spots within $S_1$ region. For each spot $s_j \in S_1$, the median gene expression across all genes is calculated, based on which $S_1$ is reordered ascendingly as $\tilde{S}_1$. Next, we simulate $N_1$ observations from $NB(\mu, s)$, which are sorted in ascending order and assigned to correspondingly ranked spots in $\tilde{S}_1$. This ordered assignment helps to preserve the typical spatial gene expression structure inherent in the ST dataset. Thereby, we obtain a simulated gene expressed at the top $t\%$ quantile level within the $S_1$ region. We can repeat this procedure to make the simulated gene to be expressed at specified quantile levels within arbitrary tissue regions.

**SpaCEX-ETC.** As shown in *Main Text*, Fig. 7*A*, SpaCEX-ETC is essentially a GAN model consisting of a generator $\boldsymbol{\mathcal{G}}$ and a discriminator $\boldsymbol{\mathcal{D}}$. Specifically, let $Y \in R^{N \times H}$ denote the original gene expression matrix from a dataset with full transcriptomic coverage, where $N$ is the number of genes and $H$ is the number of spatial spots within the dataset. The encoder of SpaCEX's MAE is denoted as $E$ with parameters $W$. Then, we have SEGs matrix as $S \in R^{N \times D} = E(Y, W)$, where $D$ is the SGE dimension. $S$ serves as the inputs to SpaCEX-ETC's generator. Let $X \in R^{\hat{N} \times M}$ denote the gene expression matrix of the target FISH-based dataset, where $\hat{N} \ll N$ represents the number of its covered genes, while $M$ represents the number of spatial spots. SpaCEX-ETC's generator consists of three subcomponents: an encoder, a decoder, and a memory bank. Explicitly, for a given gene $i$ included in $X$, the encoder passes its SEG, $s_i$, through several feed-forward layers to generate a non-linear projection $\boldsymbol{z}_i \in \boldsymbol{R}^{\boldsymbol{d}}$. The memory bank is an embedding queue $\boldsymbol{Q} \in \mathbb{R}^{N_{mem} \times d}$ filled with $\boldsymbol{z}$, where $N_{mem}$ denotes the number of in-memory embeddings. It provides an attention-based means to reconstruct $\boldsymbol{z}$ as $\tilde{\boldsymbol{z}} \in \mathbb{R}^d$:

$$\tilde{\boldsymbol{z}}_i = \boldsymbol{Q}^T \text{softmax}\left(\frac{\boldsymbol{Q}\boldsymbol{z}_i}{\tau}\right),$$

where $\tau$ is a temperature hyperparameter. $\boldsymbol{Q}$ is continuously updated during training by enqueuing recent $\tilde{\boldsymbol{z}}$ and dequeuing the oldest. This updating strategy maintains a balance between preserving previously learnt features and adapting to new features to mitigate the mode collapse risk. The decoder is a Multi-Layer Perceptron (MLP) network for regenerating the spatial gene expression vector $\hat{\boldsymbol{x}}_i$ from $\tilde{\boldsymbol{z}}_i$. SpaCEX-ETC's discriminator $\boldsymbol{\mathcal{D}}$ consists of an MLP-based encoder and a classifier. It is trained to distinguish between $\boldsymbol{x}_i \in X$ and the corresponding $\hat{\boldsymbol{x}}_i$. The total loss functions for the generator ($\mathcal{L}_{Gen}$) the discriminator ($\mathcal{L}_D$) are defined as:

$$\hat{\boldsymbol{x}} = \boldsymbol{\mathcal{G}}(s),$$
$$\mathcal{L}_{Gen} = \alpha\mathcal{L}_{rec} + \beta\mathcal{L}_{adv} = \alpha\mathbb{E}(\|\boldsymbol{x} - \hat{\boldsymbol{x}}\|_1) - \beta(\mathbb{E}[\boldsymbol{\mathcal{D}}(\hat{\boldsymbol{x}})]),$$
$$\mathcal{L}_D = \mathbb{E}[\boldsymbol{\mathcal{D}}(\hat{\boldsymbol{x}})] - \mathbb{E}[\boldsymbol{\mathcal{D}}(x)] + \lambda\mathbb{E}[(\|\nabla\boldsymbol{\mathcal{D}}(\xi)\|_2 - 1)^2],$$

where $\xi = \epsilon\hat{x} + (1-\epsilon)x, \epsilon \in (0,1)$. Here, $\mathcal{L}_{rec}$ denotes the gene reconstruction loss, while $\mathcal{L}_{adv}$ the adversarial loss. $\alpha, \beta, and\, \lambda \geq 0$ represent the weights of each loss function. $\boldsymbol{\mathcal{D}}(\hat{\boldsymbol{x}}_i) \in \boldsymbol{R}^{\boldsymbol{h}}$ is the discriminator's output for $\hat{\boldsymbol{x}}_i$, and $\mathbb{E}[\boldsymbol{\mathcal{D}}(\hat{\boldsymbol{x}}_i)] \in [0, 1]$ represents the probability that $\hat{\boldsymbol{x}}_i$ is classified as real by $D$. Additionally, a gradient penalty term on $\xi$ is included in $\mathcal{L}_D$ to ensure the Lipschitz continuity of the discriminator for maintaining the stability of the adversarial training process(7). The loss of SpaCEX-ETC's generator is instrumental in finetuning the weights of the MAE encoder in SpaCEX's module I, which enables the SEGs to adapt to the particular semantics inherent in the FISH-based ST dataset. This finetuning facilitates the generation of those genes uncovered in the FISH-based dataset with optimized fidelity, with its process explicitly described as:

$$W \leftarrow W - \gamma \frac{\partial \mathrm{E}(Y, W)}{\partial W} \frac{\partial \mathcal{L}_{Gen}}{\partial \mathrm{E}(Y, W)},$$

where $\gamma$ is the learning rate. Upon training completion, the spatial gene expression profile of any uncovered gene $j$ can be imputed as $\hat{\boldsymbol{x}}_j = \boldsymbol{\mathcal{G}}(s_j)$.

**SpaCEX-SVG.** For each real gene in the ST dataset, we first simulate five spatially homogeneous genes with independently and identically distributed expression profiles that mirror the real gene at each spot. Here, we assume the read counts of a given real gene at every spatial spot follows either a negative binomial (NB) or a zero-inflated negative binomial (ZINB) distribution. The former distribution has four parameters, including the mean $\mu$, the variance $v$, the dispersion $s$, and the probability of a positive event $p$, while the latter has an additional parameter, the proportion of zero inflation $\pi$. These parameters can be estimated from the observed spatial expression vector $x \in R^d$, where $d$ denotes the number of spatial spots. With the estimated parameters, either the "rnegbin" of the R package "MASS" or the "zinb" function of the R package "rzinb" are used to simulate spatially homogeneous genes that follow NB or ZINB distribution, respectively. Gene specific NB parameters are estimated by minimizing the negative log-likelihood of observed gene read counts w.r.t the parameters using PyTorch's autograd(4). The estimation of gene specific ZINB parameters is more involved. Explicitly, we have the following equations among the ZINB parameters:

$$v = \mu + \frac{\mu^2}{s},$$

$$\mu = \frac{s(1-p)}{p}.$$

The relationships between the ZINB parameters and the observed values of zero ratio r, the sample mean $m$ and the variance $\sigma^2$ of x are:

$$r = \pi + (1-\pi)p^s = \frac{\sum_i^d \mathrm{II}(x_i = 0)}{d}$$

$$m = (1-\pi)\mu = \frac{\sum_i^d x_i}{d}$$

$$\sigma^2 = \pi m^2 + (1-\pi)\mathrm{E}\left(\sum_i^d (x_i - m)^2\right) = \pi m^2 + (1-\pi)(v + \mu^2 - 2m\mu) = (1-\pi)v + \pi(1-\pi)\mu^2$$

$$= \frac{\sum_i^d (x_i - m)^2}{d}$$

We design a numerical algorithm to iteratively solve the equations and approximate the values of the ZINB parameters, as elaborated in Table S4. Finally, simulated genes are encoded into SGEs. The spatial variability score of any given real gene is calculated as the average scaled cosine dissimilarity between its SGE and those of the corresponding simulated genes. By ranking spatial variability scores in descending order, we can select a predetermined number of top-ranked genes as SVGs.

**SpaCEX-ISC.** SpaCEX-ISC requires a gene identity matrix $\mathcal{G} \in R^{N \times L}$ to indicate genes' identities in functional groups, where $N$ denotes the number of genes and $L$ the number of groups. Such a matrix can be constructed from SpaCEX-generated gene groups:

$$\mathcal{G}_{i,j} = \begin{cases} 1, if\ I(i) = j \\ 0, otherwise \end{cases}$$

where $I(i)$ is a function indicating gene $i$'s group identity. Alternatively, $\mathcal{G}$ can be constructed from a gene-gene similarity matrix, $\mathcal{S} \in R^{N \times N}$ , calculated as the normalized Gram matrix of SEGs:

$$\mathcal{S} = Diag(|Z|)^{-1} ZZ^T Diag(|Z|)^{-1},$$

where $Z \in R^{N \times D}$ denotes the SGE matrix, $N$ the number of genes, and D the SGE dimension. $\mathcal{S}$ is then converted to degree-normalized mutual $k$-nearest neighbors adjacency matrix $\tilde{\mathcal{A}} \in R^{N \times N}$:

$$\mathcal{A}_{i,j} = \begin{cases} 0, if\ i = j \\ 1, if\ i \neq j, i \in N_j(k), j \in N_i(k) \\ 0, otherwise \end{cases}$$

$$\tilde{\mathcal{A}} = \Delta^{-1} \mathcal{A},$$

where $N_i(k)$ denote the $k$-nearest neighbors of gene $i$ on $\mathcal{S}$. $\Delta$ is the diagonal degree matrix of $\mathcal{A}$, with $\Delta_{i,i} = \sum_j \mathcal{A}_{i,j}$. Shi-Malik spectral clustering(8) is performed on $\tilde{\mathcal{A}}$ to get $\mathcal{G}$. SpaCEX-SVG is then utilized to acquire gene spatial variability scores $\mathcal{V} \in R^N$, based on which a redundancy filtering matrix $\mathcal{F} \in R^{N \times N}$ is constructed:

$$\mathcal{F}_{i,i} = \begin{cases} 1, if\ R(\mathcal{V}_i) \leq \sigma \\ 0, otherwise \end{cases},$$

where $R(\mathcal{V}_i)$ is a function that returns the ranking percentile of $\mathcal{V}_i$ in $\mathcal{V}$, $\sigma$ is a prespecified threshold for controlling redundancy. Finally, $\mathcal{F}$ is applied to the original spatial gene expression matrix $X \in R^{M \times N}$, where $M$ denotes the number of spatial spots:

$$\tilde{X} = X\mathcal{F}.$$

Note that $\tilde{X}$ represents an informational-efficient version of $X$ and are subsequently fed into a two-layer convolutional graph neural network to generate spot embeddings, on which clustering is performed.

**Experimental Settings.**
***Identifying groups of spatially co-expressed genes***

Two important settings in this experiment include the target number of clusters and how to choose gene clusters for visualizing the spatial expression patterns of their member genes. The target number of gene clusters is based on the total number of genes in the dataset, aiming for an average of ~30 genes per cluster to approximate the average number of genes in a typical KEGG pathway(9). For the second setting, gene clusters are first categorized into three groups according to their intra-cluster gene similarity quantified by averaging the spatial autocorrelation measure (SAM, see "*Evaluation metrics*" in Methods) values among intra-cluster genes: the high-quality group (top third quantile), the low-quality group (bottom third quantile), and the medium-quality group (all remaining clusters). Subsequently, one cluster is randomly selected for visualization from each of the high- and medium-quality groups of the 10x-hDLPFC-151673 and ssq-mHippo datasets.

***Enrichment analysis***

Enrichment analysis is employed to identify biological functions or processes enriched in the investigated gene set. Here, we conduct GO pathway enrichment analyses on gene groups to assess their biological relevance to breast cancer. Among the three data types in the GO database (GOBPs, molecular functions and cellular components), we only consider GOBPs since they better reflect high-level functions of the investigated system. The R package clusterProfiler (v4.2.2) is used to identify GOBPs pathways enriched in the genes of the two groups. In each enrichment analysis, we report P-values with false discovery control adjusted by Benjamini-Hochberg procedure. The 20 most significantly enriched GOBPs/KEGG pathways are selected for further biological analyses.

***Cofunction analysis of intra-cluster genes***

In this study, we utilize the neighbor-voting method proposed by Ballouz et al(10) to assess the functional coherence of a gene set within a GOBP. Basically, under the assumption that genes within the same cluster are more functionally coherent and given the GOBP involvement of some genes in a gene cluster, it should yield more accurate prediction on the GOBP involvement of other genes in the same cluster. Specifically, let $d$ denote the number of genes, $q$ denote the number of GOBPs/pathways, and $\mathbf{A} \in \mathbb{R}^{d\times q}$ denote the gene annotation matrix. Here,

$$\mathbf{A}_{i,j} = \begin{cases} 1, \text{if gene i } \in \text{ the } j-\text{th GOBP} \\ 0, \text{otherwise} \end{cases}$$

Next, the scaled cosine gene-gene similarity matrix $\mathbf{S} \in \mathrm{R}^{d\times d}$ is calculated based on the gene embeddings $\mathrm{Z} = \{z_i, \forall i \in [1, d]\}$ generated by SpaCEX:

$$\mathbf{S}_{\mathrm{i,j}} = \frac{z_i^T z_j}{\gamma \|z_i\| \|z_j\|}, \quad (2)$$

where $\gamma$ is a temperature hyperparameter. The accuracy of GOBP predictions is evaluated using a three-fold cross-validation approach. In each fold, a random selection of one-third of the genes serves as the testing dataset, with their associated GOBPs masked by setting the corresponding rows in $\mathbf{A}$ to zero. For a gene $t$ in the testing dataset, the probability of its involvement in the $j$-th GOBP is calculated as:

$$p_{tj} = \sum_{i=1}^{d} \mathbf{S}_{t,i}\, \mathbf{A}_{i,j} / \sum_{i=1}^{d} \mathbf{S}_{t,i}$$

The prediction results are evaluated by examining the top 20 enriched GOBPs, using AUC metric. A higher AUC value indicates greater functional coherence among the tested genes for that GOBP. Genes involved in the selected GOBPs are obtained using R package AnnotationDbi (v1.56.2). The R package EGAD (v1.22.0) is employed to conduct the neighbor-voting-based prediction.

*Evaluating SpaCEX-generated gene embeddings*

This study comprises three analyses designed to assess the relational semantics of SGEs and their utility in gene functional ontology-related tasks. SGEs are generated from the 10x-hDLPFC-151676 dataset. In the first analysis, we collect 64-dimensional SGEs for the 26 members from the KRT-II, HLA-I and HLA-II gene families (Table S5). Subsequently, average-linkage hierarchical clustering is applied to these embeddings so that those with higher SGE correlations are positioned in closer proximity within the hierarchy. In the second analysis, gene clusters are generated by applying Leiden to a gene-gene similarity matrix computed using either SGEs or the original spatial gene expressions across eight resolution levels (5, 10, 15, 20, 25, 30, 35, and 40). Pathway enrichment analysis against the Reactome database is conducted on the gene clusters containing more than five members. For a given resolution level, we calculate gene clusters' average number of significantly enriched pathways, or high-confidence "pathways hits", with a Bonferroni-corrected P-value below 0.05(11). This average serves as an indicator of the effectiveness of gene representations—whether they are SGEs or original spatial gene expression profiles—in capturing intricate gene-gene connections. In the third analysis, an adapted GGIPN, which is essentially a multilayer perceptron classifier with residual connections, is trained to use gene embeddings as inputs for predicting the existence of interactions between gene pairs(3). The process of defining and selecting pairs of interacting (positive) and noninteracting (negative) genes to form the training and testing datasets is elaborated in Supplementary Note 1.4. Four types of embeddings are involved in this analysis, including the SEGs from the 10x-hDLPFC-151676 dataset, Gene2vec embeddings, scBERT embeddings, and randomly generated 64-dimensional embeddings that serve to determine the GGIPNN's baseline performance.

**Evaluation metrics**

*Gene clustering.* Davies-Bouldin (DB) index(12) is used to assess the performance of gene clustering:

$$\mathrm{DB} = \frac{1}{n}\sum_{i=1}^{n}\max_{i\neq j}\frac{d_i + d_j}{d_{(i,j)}}$$

$$d_i = \frac{2}{n_i(n_i-1)}\sum_{p,q\in\mathcal{G}_i, p\neq q}\delta_{p,q}$$

$$d_{(i,j)} = \delta_{c_i,c_j}$$

where $\delta_{p,q}$ denotes the distance between genes $p$ and $q$, n denotes the number of clusters, $d_i$ is the intra-cluster distance of cluster $i$ (calculated as the average distance between genes of cluster $i$), and $d_{(i,j)}$ is the inter-cluster distance between clusters $i$ and $j$ (calculated as the distance between cluster centroids $c_i$ $and$ $c_j$). DB index represents the ratio of intra-cluster distance (compactness) to inter-cluster distance (separateness), so a smaller value of DB index indicates better clustering performance. To evaluate the co-expression and the spatial coherence of gene clusters, Pearson distance and Euclidean distance are used in the calculation of DB index, respectively(13):

$$\delta_{p,q}^{pearson} = 1 - \rho_{p,q}$$

$$\delta_{p,q}^{euclid} = \sqrt{\left(x_p - x_q\right)^T W\left(x_p - x_q\right)}, W \in R^{N_x N_y \times N_x N_y},$$

where $\rho_{p,q}$ represents the Pearson correlation between genes $p$ and $q$. $N_x$ and $N_y$ denote the number of spatial spots along the horizontal and vertical axes of the spatial map, respectively. $x_p$ is the flattened expression matrix of gene $p$, and $W$ is the weight matrix calculated based on the spatial spots' locations using a Gaussian kernel, reflecting the spatial closeness of the spatial spots.

*Intra-cluster gene similarity.* Given any pair of genes within a cluster, the SAM metric, originally designed for measuring imagery similarity, is used to measure the similarity of their spatial expression patterns. Let $x$,$y \in R^n$ denote the two genes' spatial expression vectors, respectively. Their SAM value is computed as:

$$\mathrm{SAM}(\mathrm{x}, y) = \cos^{-1}\left(\frac{\sum_{i=1}^{n} x_i \cdot y_i}{\parallel x \parallel \cdot \parallel y \parallel}\right),$$

where $n$ denotes the number of spots. Then, the intra-cluster gene similarity of cluster $c_k$ is calculated as:

$$\mathrm{SAM}(c_k) = \frac{\sum_{i,j \in c_k, i \neq j} SAM\left(x_i, x_j\right)}{|c_k|},$$

where $|c_k|$ represents the number of gene pairs in $c_k$.

*Spatial variability of gene expressions*. Moran's *I* and Geary's *C*, which measure the spatial autocorrelation of a variable from different perspectives, are used to evaluate the spatial variability in gene expressions. The two metrics calculated on a gene $x$ are as follows:

$$\text{Moran's } I = \frac{\mathrm{n}}{\mathrm{w}} \frac{\sum_{\mathrm{i}=1}^{\mathrm{n}} \sum_{\mathrm{j}=1}^{\mathrm{n}} [\mathrm{w_{ij}}(\mathrm{x_i} - \bar{\mathrm{x}})(\mathrm{x_j} - \bar{\mathrm{x}})]}{\sum_{\mathrm{i}=1}^{\mathrm{n}} (\mathrm{x_i} - \bar{\mathrm{x}})^2}$$

$$\text{Geary's } C = \frac{\mathrm{n}}{2\mathrm{w}} \frac{\sum_{\mathrm{i}=1}^{\mathrm{n}} \sum_{\mathrm{j}=1}^{\mathrm{n}} [\mathrm{w_{ij}}(\mathrm{x_i} - \mathrm{x_j})^2]}{\sum_{\mathrm{i}=1}^{\mathrm{n}} (\mathrm{x_i} - \bar{\mathrm{x}})^2}$$

$$\mathrm{w_{ij}} = \begin{cases} 1, if j \in N_k(i) \\ 0, otherwise \end{cases}$$

$$\mathrm{w} = \sum_{\mathrm{i}=1}^{\mathrm{n}} \sum_{\mathrm{j}=1}^{\mathrm{n}} \mathrm{w_{ij}}$$

where $x_i$ denote the read counts at spot $I$, $\bar{x}$ the average read counts over all spots, $n$ the total number of spots, $w_{ij}$ the spatial weight between spots *i* and $j$, $N_k(i)$ the set of *k*-nearest neighbors of spot *i* in Euclidean distance. Moran's I ranges from [-1,1] and Geary's C ranges from [0,2]. Values closer to the upper boundary for Moran's I (or the lower boundary for Geary's C) indicate a stronger positive autocorrelation and a more regular spatial pattern.

*Spatial clustering.* The accuracy of clustering for spatial spots is assessed using the Adjusted Rand Index (ARI) and Normalized Mutual Information (NMI). For ARI, let $n$ denote the total number of spots, $n_{ij}$ the number of spots of domain type $i$ in cluster $j$, $a_i$ the total number of spots of domain type $i$, $b_j$ the number of all spots in cluster $j$.

$$\mathrm{ARI} = \frac{\sum_{ij}\binom{n_{ij}}{2} - \left[\sum_i \binom{a_i}{2}\sum_j \binom{b_j}{2}\right] / \binom{n}{2}}{\frac{1}{2}\left[\sum_i \binom{a_i}{2} + \sum_j \binom{b_j}{2}\right] - \left[\sum_i \binom{a_i}{2}\sum_j \binom{b_j}{2}\right] / \binom{n}{2}}$$

For NMI, let $L$ denote the true labels of spots, $\tilde{L}$ the cluster labels of spots, $MI$ the mutual information function, and H the entropy function.

$$\mathrm{NMI} = \frac{2 \times \mathrm{MI}(\tilde{\mathrm{L}}, \mathrm{L})}{\mathrm{H}(\tilde{L}) + \mathrm{H}(\mathrm{L})}$$

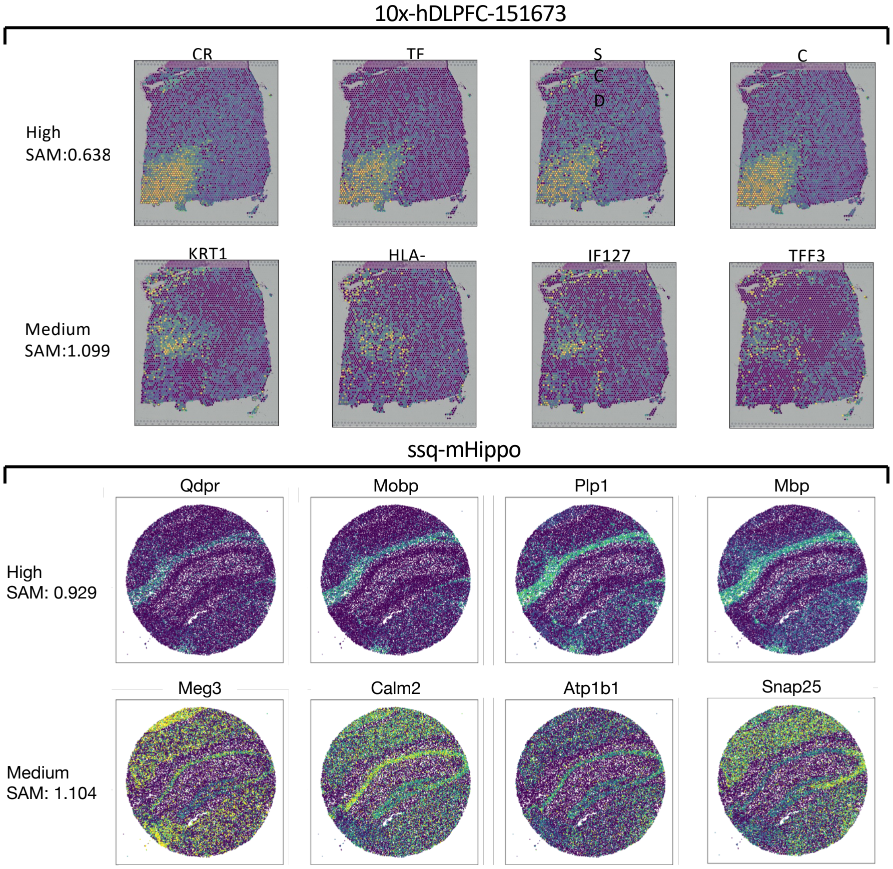


**Fig. S1.** Spatial expression patterns of individual genes within co-expression groups identified by SpaCEX from the human DLPFC 10x Visium (10x-hDLPFC-151673) and the mouse hippocampus Slide-seqV2 (ssq-mHippo) datasets, respectively. For each dataset, the denoised expression patterns of four genes randomly selected from the same group are shown in the same row. The first and second rows represent two gene clusters of high and medium intra-cluster similarity respectively, as assessed by the SAM metric. A lower SAM value indicates higher intra-cluster similarity.

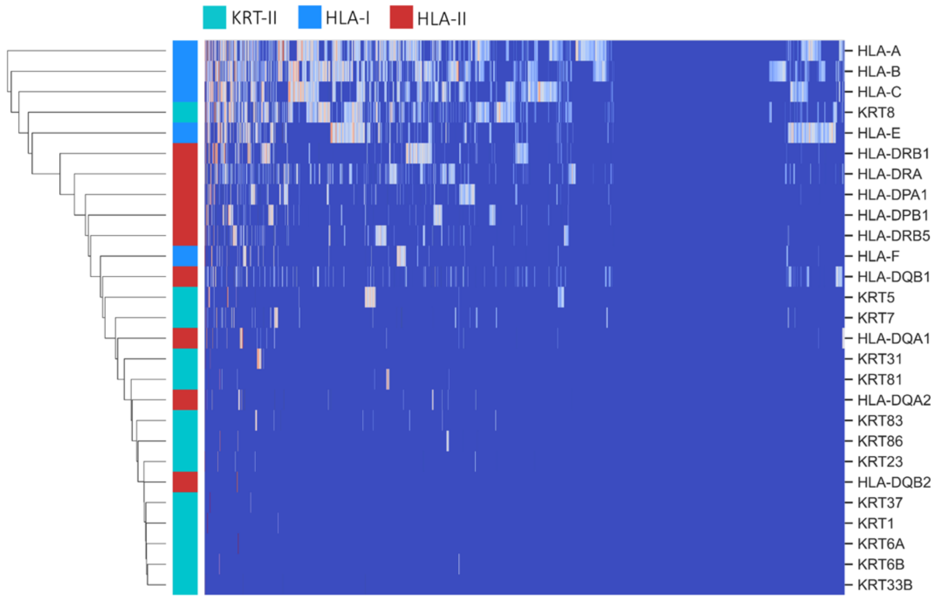


**Fig. S2.** Hierarchical clustering performed on the original spatial expression profiles of the HLA-I, HLA-II, and KRT-II gene family members in the 10x-hDLPFC-151676 dataset. Genes with correlated expression profiles are positioned in close proximity along the y-axis. The dimensions of the gene expression vectors are represented along the x-axis. Gene families are indicated in different colors on the y-axis. The color intensities in the diagram positively correlate with the magnitude of the expression values.

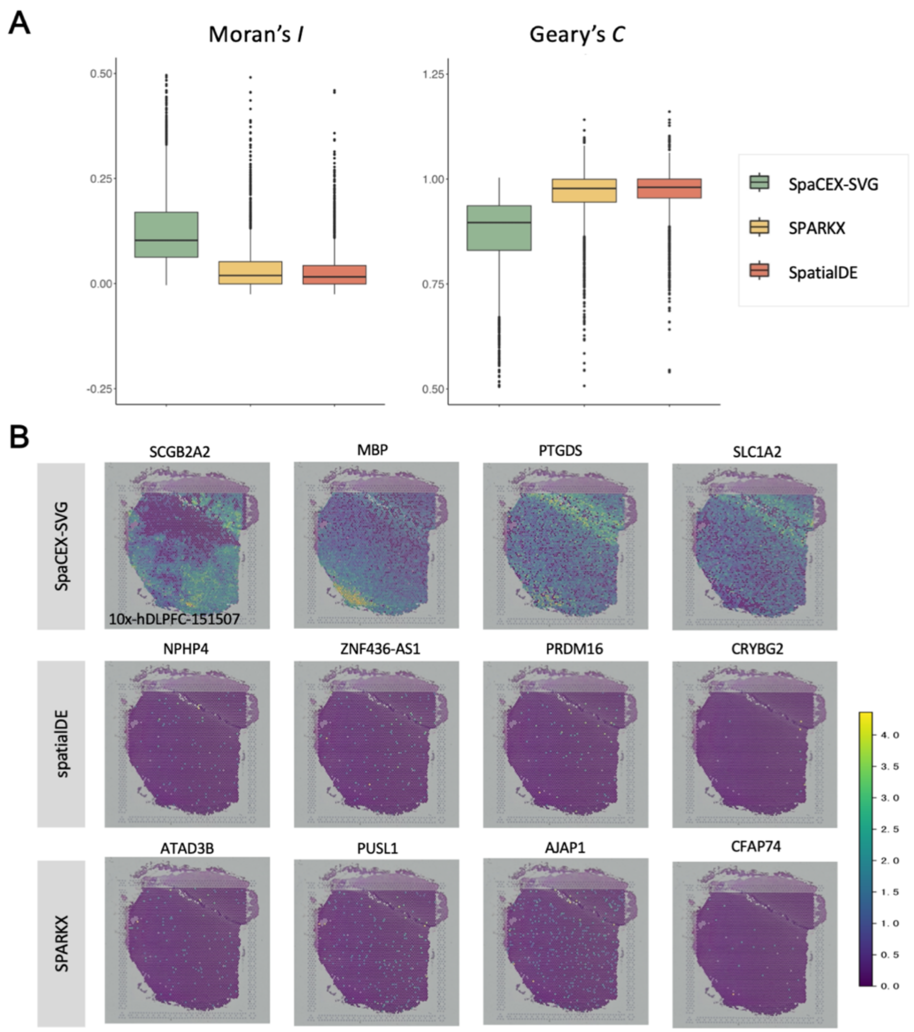


**Fig. S3.** SGE-based SVG detection with SpaCEX-SVG. (*A*) The box plots show the Moran's I (left panel) and Geary's C (right panel) indices for the top 3000 SVGs detected by SpaCEX-SVG, SPARK-X and SpatialDE from the 10x-hDLPFC-151507 dataset. The value ranges for these indices are [-1,1] for Moran's I and [0, 2] for Geary's C. A value approaching the upper limit for Moran's I (or the lower limit for Geary's C) indicate a stronger positive autocorrelation and a more pronounced spatial pattern. (*B*) The spatial expression patterns of the top four SVGs, as identified by SpaCEX-SVG, SPARK-X and SpatialDE, are displayed in the top, middle and bottom rows, respectively. The panels in the top row have the clear spatial patterns, while those in the other two rows are almost devoid of noticeable spatial patterns.

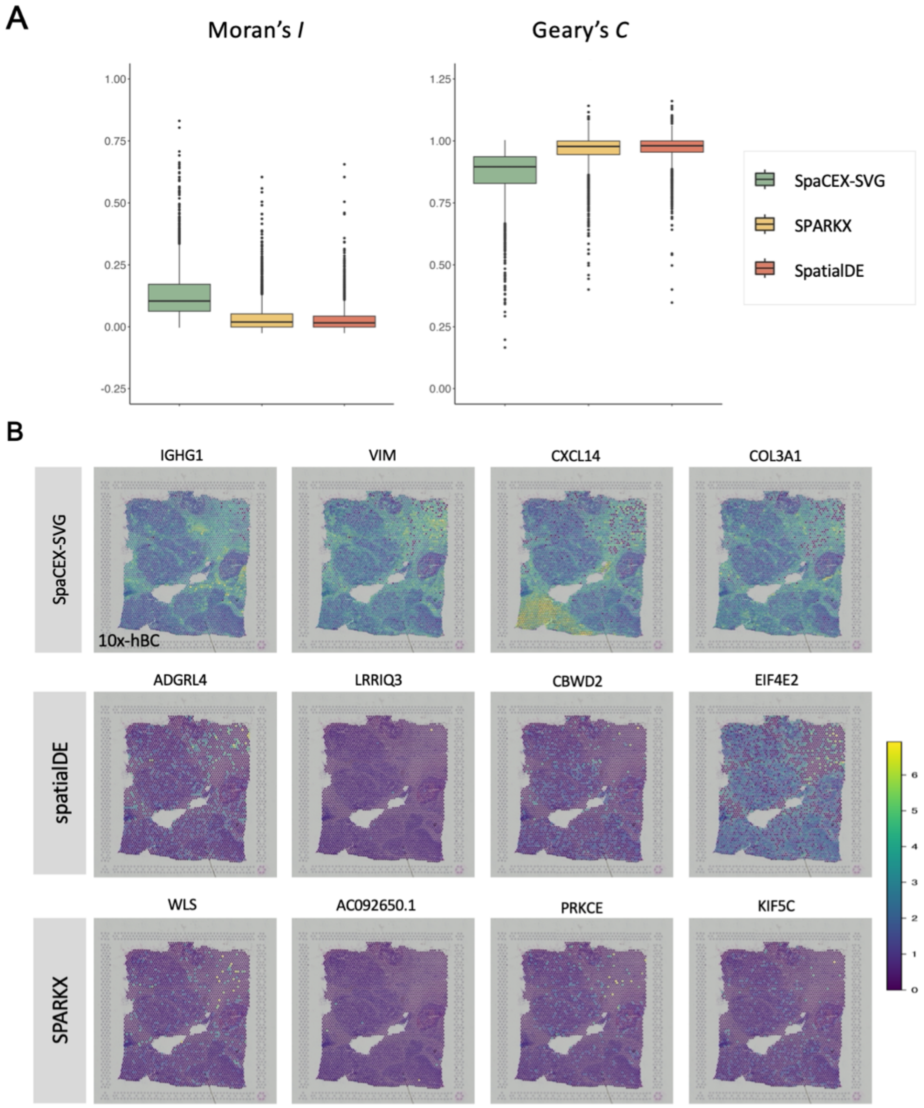


**Fig. S4.** SGE-based SVG detection with SpaCEX-SVG. (*A*), The box plots show the Moran's I (left panel) and Geary's C (right panel) indices for the top 3000 SVGs detected by SpaCEX-SVG, SPARKX and SpatialDE from the 10x-hBC dataset. The value ranges for these indices are [-1,1] for Moran's I and [0, 2] for Geary's C. A value approaching the upper limit for Moran's I (or the lower limit for Geary's C) indicate a stronger positive autocorrelation and a more pronounced spatial pattern. (*B*), The spatial expression patterns of the top four SVGs, as identified by SpaCEX-SVG, SPARK-X, and SpatialDE, are displayed in the top, middle and bottom rows, respectively. The panels in the top row exhibit the clearest spatial patterns among all others.

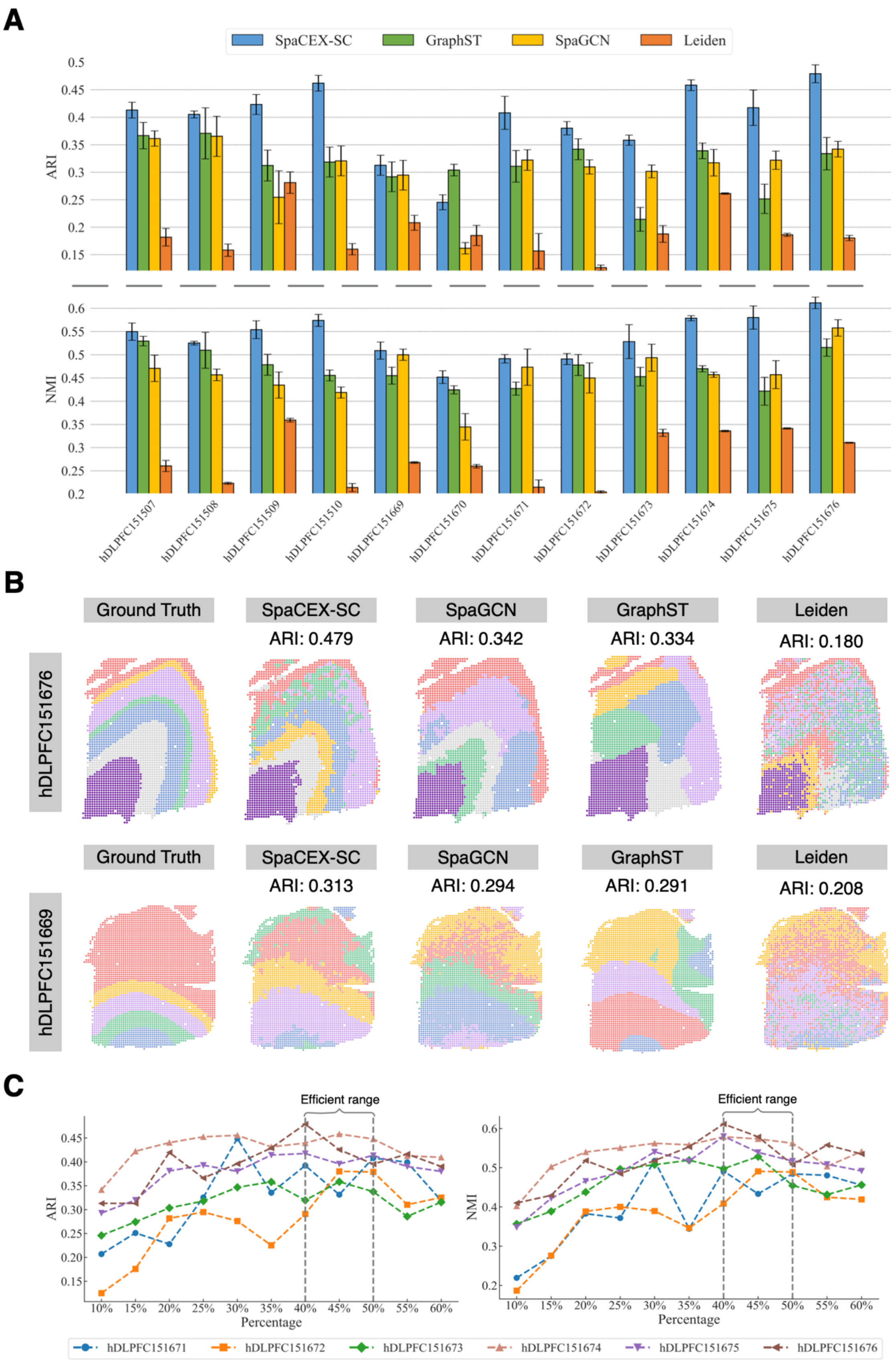
A
SpaCEX-SC
GraphST
SpaGCN
Leiden
ARI
NMI
hDLPFC151507
hDLPFC151508
hDLPFC151509
hDLPFC151510
hDLPFC151669
hDLPFC151670
hDLPFC151671
hDLPFC151672
hDLPFC151673
hDLPFC151674
hDLPFC151675
hDLPFC151676
B
Ground Truth
SpaCEX-SC
SpaGCN
GraphST
Leiden
hDLPFC151676
ARI: 0.479
ARI: 0.342
ARI: 0.334
ARI: 0.180
hDLPFC151669
ARI: 0.313
ARI: 0.294
ARI: 0.291
ARI: 0.208
C
Efficient range
ARI
NMI
Percentage
hDLPFC151671
hDLPFC151672
hDLPFC151673
hDLPFC151674
hDLPFC151675
hDLPFC151676

**Fig. S5.** SGE-based spatial clustering with SpaCEX-SC. (*A*) The bar plots illustrate the accuracy of spatial clustering across twelve 10x-hDLPFC datasets using SpaCEX-SC, GraphST, SpaGCN and Leiden in terms of ARI in the first row and NMI in the second row. (*B*) We randomly select two datasets (10x-hDLPFC-151676 and 10x-hDLPFC-151669) to visualize their ground truth domain annotations and spatial clustering results across methods. (*C*) The line plots showcase the trend of spatial clustering accuracies with percentages of SpaCEX-SC-filtered redundant gene information across the six 10x-hDLPFC datasets. The spatial clustering performances peaks when approximately 50%-60% redundant information is excluded.

**Table S1. Datasets used**

| Dataset | Species | Tissue | Protocol | No. spots | No. genes | Health status | Source |
|---|---|---|---|---|---|---|---|
| ssq-mHippo | Mouse | Hippocampus | Slide-seqV2 | 41786 | 4000 | Healthy | Stickles et al(14) |
| sqf-mEmb | | Embryo tissues | SeqFish | 19416 | 351 | | Lohoff et al(15) |
| 10x-mEmb | | | 10x Visium | 487 | 16901 | | Chen et al(16) |
| 10x-hDLPFC-151507 | Human | DLPFC | | 4226 | 18400 | | Maynard et al(17) |
| 10x-hDLPFC-151508 | | | | 4384 | 18033 | | |
| 10x-hDLPFC-151509 | | | | 4789 | 18663 | | |
| 10x-hDLPFC (151510) | | | | 4634 | 18420 | | |
| 10x-hDLPFC (151669) | | | | 3661 | 18543 | | |
| 10x-hDLPFC (151670) | | | | 3498 | 18303 | | |
| 10x-hDLPFC (151671) | | | | 4110 | 19002 | | |
| 10x-hDLPFC (151672) | | | | 4015 | 18636 | | |
| 10x-hDLPFC (151673) | | | | 3639 | 19119 | | |
| 10x-hDLPFC (151674) | | | | 3673 | 19836 | | |
| 10x-hDLPFC (151675) | | | | 3592 | 18619 | | |
| 10x-hDLPFC (151676) | | | | 3460 | 18639 | | |
| 10x-hBC | | Breast | | 3798 | 22218 | Cancer | 10x Genomics: https://support.10xgenomics.com/spatial-gene-expression/datasets/1.1.0/V1_Breast_Cancer_Block_A_Section_1 |

| | | | | | | | |
|---|---|---|---|---|---|---|---|
| 10x-hMTG-2-3 | | MTG | | 4838 | 36601 | AD | Chen et al(18) |
| 10x-hMTG-1-1 | | | | 3742 | 36601 | Healthy | |
| 10x-hMTG-18-64 | | | | 4225 | 36601 | Healthy | |

**Table S2. List of benchmark methods.**

| Experiments | Methods | Basic method | Implementation | Reference |
|---|---|---|---|---|
| Co-expressed gene identification | CNN-PReg | Deep embedded clustering with graph Laplacian regularization | https://github.com/kuanglab/CNN-PReg | Song et al(13). |
| | Giotto | Hierarchical clustering on genes | R package *Giotto* 3.2 | Dries, et al(19). |
| | STUtility | Non-negative matrix factorization | R package *STUtility* 1.1.1 | Bergenstråhle et al(20). |
| | SPARK | Hierarchical clustering on genes | R package *SPARK* 1.1.1 | Sun et al(21). |
| Transcriptomic coverage enhancement | Tangram | Impute missing ST genes via single-cells mapped to spatial locations based on a probability mapping matrix | https://github.com/tangrams/tangram | Biancalani et al(22) |
| | SpaGE | Mapping single-cells to spatial location via cross-sample alignment between scRNA-seq and ST in a latent space, followed by missing gene imputation using KNN single-cells | https://github.com/tabdelaal/SpaGE | Abdelaal et al(23) |
| | SpaOTsc | Mapping single-cells to spatial locations via structured optimal transport, followed by imputing missing ST genes from scRNA-seq data | https://github.com/zcang/SpaOTsc | Cang and Nie(24) |
| SVG detection | SpatialDE | The Q-value for spatial differential expression based on Gaussian process regression | Python package *SpatialDE* 1.1.3 | Svensson et al(25). |
| | SPARKX | The adjusted P-value based on a robust covariance test framework | R package *SPARK* 1.1.1 | Zhu et al(26). |
| Spatial clustering | SpaGCN | Spatial clustering by graph convolutional network using Gene spatial expression information | https://github.com/jianhuupenn/SpaGCN | Hu et al(27). |
| | GraphST | The graph-based self-supervised contrastive learning method | https://github.com/JinmiaoChenLab/GraphST | Long et al(28). |
| | Leiden | Graph community detection | Python package *Scanpy* 1.9.6 | Traag et al(29). |
| Gene-gene interaction prediction | Gene2vec | Skip-gram-like neural network | https://github.com/jingcheng-du/Gene2vec | Du et al(3). |
| | scBERT | Bidirectional BERT-like method for generating gene embeddings based on predicting masked genes. | https://github.com/TencentAILabHealthcare/scBERT | Yang et al(30) |

**Table S3. List of AD-associated genes.**

| Number | Gene | Reference |
|---|---|---|
| 1 | APP | Levy et al(31) |
| 2 | PSEN1 | Levy-Lahad et al(32) |
| 3 | PSEN2 | Levy-Lahad et al(32) |
| 4 | ACE | Hemming and Selkoe(33) |
| 5 | TREM2 | Neumann and Daly(34) |
| 6 | SORL1 | Pottier et al(35) |
| 7 | ABCA7 | Lacour et al(36) |
| 8 | CD33 | Bertram et al(37) |
| 9 | ATXN1 | Zhang et al(38) |
| 10 | CHRNB2 | Cook et al(39) |
| 11 | CHRNA4 | Cook et al(39) |
| 12 | MAPT | Myers et al(40) |
| 13 | CST3 | Bertram and Tanzi(41) |
| 14 | TF | Robson et al(42) |
| 15 | GAB2 | Reiman et al(43) |
| 16 | APOE | Grupe et al(44) |
| 17 | ACAN | Grupe et al(44) |
| 18 | BCR | Grupe et al(44) |
| 19 | CTSS | Grupe et al(44) |
| 20 | PLD3 | Giri et al(45) |
| 21 | UNC5C | Giri et al(45) |
| 22 | AKAP9 | Giri et al(45) |
| 23 | ADAM10 | Giri et al(45) |
| 24 | CLU | Giri et al(45) |
| 25 | CR1 | Giri et al(45) |
| 26 | BIN1 | Giri et al(45) |
| 27 | CD2AP | Giri et al(45) |
| 28 | PICALM | Giri et al(45) |
| 29 | HLA-DRB5 | Giri et al(45) |
| 30 | HLA-DRB1 | Giri et al(45) |
| 31 | INPP5D | Giri et al(45) |
| 32 | MEF2C | Giri et al(45) |
| 33 | CASS4 | Giri et al(45) |
| 34 | PTK2B | Giri et al(45) |

| | | |
|---|---|---|
| 35 | ZCWPW1 | Giri et al(45) |
| 36 | CELF1 | Giri et al(45) |
| 37 | FERMT2 | Giri et al(45) |
| 38 | SLC24A4 | Giri et al(45) |
| 39 | RIN3 | Giri et al(45) |
| 40 | IQCK | Kunkle et al(46) |
| 41 | ADAMTS1 | Kunkle et al(46) |
| 42 | WWOX | Kunkle et al(46) |

**Table S4. Estimating ZINB parameters given the spatial expressions of a gene**

**Input**: target gene's expression vector: $x \in R^d$; max iteration: MaxIter; stopping threshold: $\delta$;

**Output**: $\pi$; $\mu$; $v$; $s$; $p$;

$r = \frac{\sum_i^d II(x_i=0)}{d}, m = \frac{\sum_i^d x_i}{d}, \sigma^2 = \frac{\sum_i^d (x_i - m)^2}{d}$

$\pi_0 = \alpha \cdot r, \alpha \in (0,1)$. Due to the sparsity of gene count matrix, $\alpha$ is set to be 0.99 in our case.

**for** $j \in [0, \text{MaxIter}]$ ***do***

$\quad \mu_j = \frac{m}{1-\pi_j}$

$\quad v_j = \max\left(\frac{\sigma^2 - \pi_j(1-\pi_j)\,\mu_j^2}{1-\pi_j}, 1e-5\right)$

$\quad s_j = \max\left(\frac{\mu_j^2}{v_j - \mu_j}, 1\right)$

$\quad p_j = \max\left(\frac{s_j}{\mu_j + s_j}, 1e-6\right)$

$\quad \pi_{j+1} = \frac{r - p_j^{s_j}}{1 - p_j^{s_j}}$

$\quad$ **if** $|\pi_j - \pi_{j+1}| \leq \delta$, **then** Stop

**end for**

**return** $\pi = \pi_{j+1}$; $\mu = \mu_j$; $v = v_j$; $s = s_j$; $p = p_j$

**Table S5. List of members of three gene families involved in SGE-based hierarchical clustering**

| Number | Gene | Gene family |
|---|---|---|
| 1 | KRT1 | KRT-II |
| 2 | KRT5 | KRT-II |
| 3 | KRT6A | KRT-II |
| 4 | KRT6B | KRT-II |
| 5 | KRT7 | KRT-II |
| 6 | KRT8 | KRT-II |
| 7 | KRT23 | KRT-II |
| 8 | KRT31 | KRT-II |
| 9 | KRT33B | KRT-II |
| 10 | KRT37 | KRT-II |
| 11 | KRT81 | KRT-II |
| 12 | KRT83 | KRT-II |
| 13 | KRT86 | KRT-II |
| 14 | HLA-A | HLA-I |
| 15 | HLA-B | HLA-I |
| 16 | HLA-C | HLA-I |
| 17 | HLA-E | HLA-I |
| 18 | HLA-F | HLA-I |
| 19 | HLA-DPA1 | HLA-II |
| 20 | HLA-DPB1 | HLA-II |
| 21 | HLA-DQA1 | HLA-II |
| 22 | HLA-DQA2 | HLA-II |
| 23 | HLA-DQB1 | HLA-II |
| 24 | HLA-DQB2 | HLA-II |
| 25 | HLA-DRA | HLA-II |
| 26 | HLA-DRB1 | HLA-II |
| 27 | HLA-DRB5 | HLA-II |

**Table S6. List of the 20 most significantly enriched GOBPs in the three gene clusters generated by SpaCEX and in the control cluster.**

| Cluster | GOBP ID | GOBP name | Related cancer cell type |
|---|---|---|---|
| C1 (IDC) | GO:0006986 | response to unfolded protein | Invasive cancer |
| | GO:0035966 | response to topologically incorrect protein | Invasive cancer |
| | GO:0071353 | cellular response to interleukin-4 | Invasive cancer |
| | GO:0070670 | response to interleukin-4 | Invasive cancer |
| | GO:0045793 | positive regulation of cell size | Cancer |
| | GO:0034605 | cellular response to heat | Cancer |
| | GO:0043627 | response to estrogen | Invasive cancer |
| | GO:1902947 | regulation of tau-protein kinase activity | Invasive cancer |
| | GO:0033599 | regulation of mammary gland epithelial cell proliferation | Cancer |
| | GO:0006457 | protein folding | Invasive cancer |
| | GO:0007163 | establishment or maintenance of cell polarity | Benign stroma |
| | GO:0034620 | cellular response to unfolded protein | Invasive cancer |
| | GO:0044321 | response to leptin | Invasive cancer |
| | GO:0051131 | chaperone-mediated protein complex assembly | Invasive cancer |
| | GO:0006123 | mitochondrial electron transport, cytochrome c to oxygen | Others |
| | GO:0035162 | embryonic hemopoiesis | Others |
| | GO:0009408 | response to heat | Cancer |
| | GO:0033598 | mammary gland epithelial cell proliferation | Cancer |
| | GO:0035967 | cellular response to topologically incorrect protein | Invasive cancer |
| | GO:0051897 | positive regulation of protein kinase B signaling | Invasive cancer |
| C2 (DCIS) | GO:0002181 | cytoplasmic translation | Others |
| | GO:0030277 | maintenance of gastrointestinal epithelium | Invasive cancer |
| | GO:0006096 | glycolytic process | Invasive cancer |
| | GO:0006757 | ATP generation from ADP | Cancer |
| | GO:0046031 | ADP metabolic process | Cancer |
| | GO:0010669 | epithelial structure maintenance | Invasive cancer |
| | GO:0006165 | nucleoside diphosphate phosphorylation | Cancer |
| | GO:0046939 | nucleotide phosphorylation | Cancer |
| | GO:0009135 | purine nucleoside diphosphate metabolic process | Cancer |
| | GO:0009179 | purine ribonucleoside diphosphate metabolic process | Cancer |

| | | | |
|---|---|---|---|
| | GO:0006090 | pyruvate metabolic process | Invasive cancer |
| | GO:0009185 | ribonucleoside diphosphate metabolic process | Cancer |
| | GO:0009132 | nucleoside diphosphate metabolic process | Others |
| | GO:0030879 | mammary gland development | Invasive cancer |
| | GO:0016052 | carbohydrate catabolic process | Invasive cancer |
| | GO:0006006 | glucose metabolic process | Invasive cancer |
| | GO:0042273 | ribosomal large subunit biogenesis | Cancer |
| | GO:0006735 | NADH regeneration | Invasive cancer |
| | GO:0061621 | canonical glycolysis | Cancer |
| | GO:0061718 | glucose catabolic process to pyruvate | Cancer |
| **C3 (benign stroma)** | GO:0006958 | complement activation, classical pathway | Invasive cancer |
| | GO:0002455 | humoral immune response mediated by circulating immunoglobulin | Invasive cancer |
| | GO:0006956 | complement activation | Invasive cancer |
| | GO:0016064 | immunoglobulin mediated immune response | Benign stroma |
| | GO:0019724 | B cell mediated immunity | Benign stroma |
| | GO:0002449 | lymphocyte mediated immunity | Benign stroma |
| | GO:0002460 | adaptive immune response based on somatic recombination of immune receptors built from immunoglobulin superfamily domains | Benign stroma |
| | GO:0006911 | phagocytosis, engulfment | Invasive cancer |
| | GO:0099024 | plasma membrane invagination | Benign stroma |
| | GO:0006959 | humoral immune response | Others |
| | GO:0010324 | membrane invagination | Cancer |
| | GO:0002443 | leukocyte mediated immunity | Benign stroma |
| | GO:0050871 | positive regulation of B cell activation | Benign stroma |
| | GO:0006910 | phagocytosis, recognition | Cancer |
| | GO:0002253 | activation of immune response | Cancer |
| | GO:0050864 | regulation of B cell activation | Benign stroma |
| | GO:0050853 | B cell receptor signaling pathway | Benign stroma |
| | GO:0042113 | B cell activation | Benign stroma |
| | GO:0051251 | positive regulation of lymphocyte activation | Benign stroma |
| | GO:0002696 | positive regulation of leukocyte activation | Benign stroma |
| **C4 (control)** | GO:0071712 | ER-associated misfolded protein catabolic process | Invasive cancer |
| | GO:0071218 | cellular response to misfolded protein | Invasive cancer |
| | GO:0051788 | response to misfolded protein | Invasive cancer |

| | | | |
|---|---|---|---|
| | GO:0006515 | protein quality control for misfolded or incompletely synthesized proteins | Invasive cancer |
| | GO:0032717 | negative regulation of interleukin-8 production | Others |
| | GO:0006383 | transcription by RNA polymerase III | Cancer |
| | GO:0098781 | ncRNA transcription | Invasive cancer |
| | GO:0016073 | snRNA metabolic process | Others |
| | GO:0055072 | iron ion homeostasis | Others |
| | GO:0051952 | regulation of amine transport | Others |
| | GO:0016575 | histone deacetylation | Cancer |
| | GO:0032677 | regulation of interleukin-8 production | Cancer |
| | GO:0032637 | interleukin-8 production | Invasive cancer |
| | GO:0015837 | amine transport | Others |
| | GO:0036503 | ERAD pathway | Others |
| | GO:0035967 | cellular response to topologically incorrect protein | Invasive cancer |
| | GO:0006476 | protein deacetylation | Others |
| | GO:0018401 | peptidyl-proline hydroxylation to 4-hydroxy-L-proline | Others |
| | GO:0033483 | gas homeostasis | Others |
| | GO:0051342 | regulation of cyclic-nucleotide phosphodiesterase activity | Others |
